# Jam2 Signaling and Hand2 Function are Required for the Formation of Organ-Specific Vascular Progenitors in Zebrafish

**DOI:** 10.1101/2025.10.10.681632

**Authors:** Martyna Griciunaite, Julius Martinkus, Sanjeeva Metikala, Ricardo DeMoya, Suman Gurung, Surya Prakash Rao Batta, Vishal Y. Mardhekar, Myles A. Hammond, Luiza N. Loges, Diandra Rufin Florat, Krishna K. Rentachintala, Saulius Sumanas

**Affiliations:** Department of Pathology and Cell Biology, University of South Florida, Tampa, FL, USA

## Abstract

The mechanisms regulating the formation of organ-specific vasculature are still poorly understood. We have previously identified a population of late-forming endothelial progenitor cells in zebrafish embryos (termed secondary vascular field, SVF), which emerge from the lateral plate mesoderm after 24 hpf stage, when blood circulation has already been initiated. Here, we investigate the functional role of SVF cells and the molecular mechanisms that govern the emergence of these SVF cells and their contribution to vasculature. We found that the bHLH transcription factor Hand2 and junctional adhesion molecule Jam2b are expressed in the SVF-forming region and are required for the emergence of SVF cells in zebrafish embryos. Time-lapse imaging and *jam2b:Cre*-based lineage tracing showed that SVF cells serve as the major source of the intestinal vasculature, including the supraintestinal artery (SIA) and subintestinal vein (SIV), which are subsequently remodeled to provide blood flow to many internal organs. To analyze the functional role of *jam2b* and the related *jam2a* gene in vascular development, we generated double maternal-zygotic *jam2a; jam2b* mutants, which display a greatly reduced number of SVF cells and show defects in the development of the intestinal vasculature. Further analysis showed that *hand2* functions in the SVF-forming region upstream of *jam2b* and is required to induce expression of the transcription factor *etv2/etsrp,* a known master regulator of vasculogenesis. In summary, our results identify new roles for Jam2 signaling and Hand2 function in the emergence of organ-specific vascular progenitors. The presence of similar progenitors in mammalian embryos suggests that this mechanism is evolutionarily conserved.

## INTRODUCTION

It has been well established that organ-specific blood vessels exhibit a large diversity of specialized endothelial cells. However, their developmental origin remains debated. Previous studies have suggested that the progenitors of internal organ vasculature in mammalian embryos form by differentiation *de novo,* or vasculogenesis.^1-4^ It has been demonstrated that the endocardium, endoderm, and mesothelium may contribute to the organ vasculature in mammals and avian embryos.^5-7^ In contrast, several studies have argued that the posterior cardinal vein (PCV) serves as the major source of organ-specific vascular endothelial cells in zebrafish embryos.^8-10^ According to this model, individual cells from the PCV migrate between 30 and 48 hours post-fertilization (hpf) and coalesce into the intestinal vessels, which include the supraintestinal artery (SIA) and the subintestinal vein (SIV). These vessels subsequently undergo remodeling and ultimately provide blood supply to the internal organs, including the pancreas and the liver.^10^ However, this model in zebrafish contrasts with the *de novo* vasculogenesis model suggested in mammalian embryos. In addition, it is unclear whether the PCV is the only or even the main source of internal organ-specific vascular progenitors.

In zebrafish, similar to mammalian embryos, the earliest vascular progenitors originate from the lateral plate mesoderm (LPM) during late gastrulation and somitogenesis, and coalesce into the major axial vessels, the dorsal aorta (DA) and the PCV.^11-14^ This process is thought to be largely complete by the 24 hpf stage, when blood circulation is first initiated. We have recently demonstrated that between 24 and 48 hpf a separate group of late-forming vascular progenitors emerges from the LPM along the yolk extension at the site termed the secondary vascular field (SVF).^15^ SVF cells can uniquely incorporate into the PCV after blood circulation has already been established, and show high expression of vascular progenitor markers, including *etv2/etsrp, tal1/scl,* and *npas4l,* but are negative for the expression of more differentiated endothelial markers such as *kdrl* or *fli1a.* Previous time-lapse imaging using *etv2* reporter lines demonstrated that SVF cells can contribute to the intestinal vasculature, including the SIA and SIV.^15^ However, the ultimate fate of SVF cells has not been clear due to the absence of genetic tools to perform lineage tracing. Furthermore, mechanisms regulating the emergence of SVF cells and their contribution to vasculature remain largely unknown.

To identify the molecular signature of SVF cells, we previously performed transcriptomic analysis of the zebrafish trunk region at 30 hpf.^16^ This analysis identified several SVF marker genes that had not previously been associated with vascular development, including *junctional adhesion molecule 2b (jam2b).*^15^ Vertebrate JAM homologs have been implicated in diverse processes including leukocyte-endothelial interaction and transendothelial migration, maintenance of hematopoietic stem cells (HSCs), and vascular development.^17,18^ A related zebrafish paralog *jam2a* has been previously implicated in HSC formation.^19^ However, the role of *jam2b* in vascular development has not been described.

Here we analyzed the role of zebrafish *jam2b* in the formation of intestinal vasculature and the emergence of SVF cells. We demonstrate that *jam2b* is expressed along the lateral plate mesoderm in the region that includes SVF cells. Using genetic lineage tracing and time-lapse imaging, we demonstrate that *jam2b-*positive cells contribute to multiple tissues, including the mesothelium, pericardium, fin buds and venous and intestinal vasculature. To study the role of Jam2 homologs in vascular development, we have generated genetic mutants in *jam2b* and a related homolog, *jam2a*. Double *jam2a; jam2b* maternal-zygotic mutants show a reduced number of SVF cells, which correlates with defects in the formation of the intestinal and venous vasculature; they also display reduced regenerative potential after endothelial cell ablation. We further demonstrate that the transcription factor *hand2* functions upstream of *jam2b* and is required for the formation of SVF cells. Altogether, our results identify a new role for Jam2 signaling and Hand2 function in the emergence of organ-specific vascular progenitors.

## RESULTS

### *Jam2b* expression analysis

In our previous transcriptomic analysis, we identified *jam2b* as one of the markers for SVF cells, which are late-forming vascular progenitors that emerge along the yolk extension between 24 and 48 hpf.^15^ As we and others have previously demonstrated, *jam2b* expression is localized to the ventrolateral region of the LPM along the anterior and trunk regions between the 20 somite and 48 hpf stages.^15,20,21^ It is bilaterally expressed along the yolk extension between 24 and 48 hpf, where SVF cells originate (Suppl. Fig. S1). We have previously demonstrated that *jam2b* expression partially overlaps with the expression of the vascular progenitor marker *etv2/etsrp* in SVF cells along the medial portion of the *jam2b* expression domain within the mid-trunk region.^15^

### Generation and characterization of *jam2b^Gt(2A-Gal4)^* knock-in reporter line

To analyze *jam2b+* cells in live embryos, we used a CRISPR / Cas9-mediated non-homologous recombination approach ^22^ to insert the Gal4 transcriptional activator into the genomic locus of *jam2b* (Suppl. Fig. S2A). Similar to the endogenous *jam2b* mRNA expression, *jam2b^Gt(2A-Gal4)ci55^;UAS:GFP* (further abbreviated as *jam2b^Gt(2A-Gal4)^;UAS:GFP*) expression was observed bilaterally in the lateral plate mesoderm along the yolk and yolk extension at 24 hpf (Suppl. Fig. S2B; Fig. 1A,B). At 3 and 5 dpf, *jam2b^Gt(2A-Gal4)^;UAS:GFP* expression was observed in the cell layer surrounding the yolk and yolk extension, which presumptively corresponds to the mesothelium (Fig. 1C-F). GFP expression was also observed in subsets of skeletal and craniofacial muscle cells, and in the cells at the periphery of the eye (Fig. 1C-F). GFP expression in the trunk portion of the LPM partially overlapped with *etv2* expression in SVF cells at 30 hpf (Fig. 1G-I). Interestingly, GFP-positive cells were observed in the vasculature between 30 hpf and 3 dpf (Fig. 1J-R). The majority of GFP+ cells were located in the PCV, while a few cells were found within the ISVs and SIA; less frequently, contribution to the DA and parachordal lymphangioblasts (PLs) was observed (quantified in Suppl. Fig. S3). In addition, some GFP-positive cells were located in the thoracic duct at 4 dpf, suggesting that SVF cells contribute to lymphatic vasculature (Fig. 1S-U). Due to very bright GFP expression over the yolk, we could not reliably analyze the presence of GFP-positive cells in the SIV, which is positioned over the yolk and yolk extension. The number of GFP-positive cells in the vasculature was greatly reduced in *etv2*-inhibited embryos, while expression of *jam2b^Gt(2A-Gal4);^ UAS:GFP* within the LPM was not significantly affected (Suppl. Fig. S4), suggesting that vascular contribution of *jam2b+* cells depends on *etv2* function. Overall, these findings suggest that *jam2b+* cells contribute to the vasculature, including the PCV, ISVs, intestinal vasculature, and lymphatics. These findings are consistent with our previously reported contribution of SVF cells to the PCV and intestinal vasculature ^15^.

**Figure 1.**
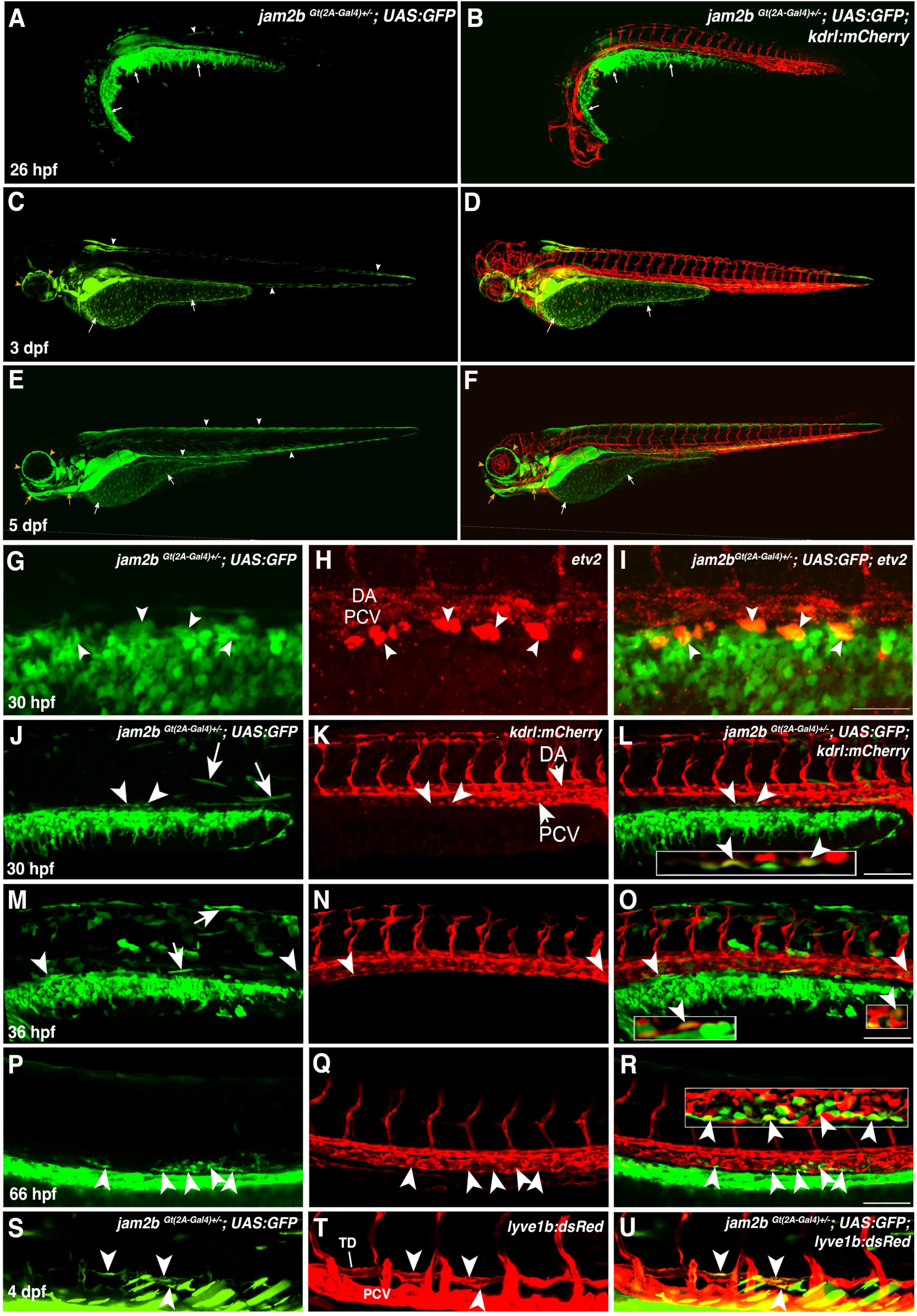
Characterization of *jam2b^Gt(2A-Gal4)^; UAS:GFP* expression at 1-5 dpf. (A-F) *jam2b^Gt(2A-Gal4)^;UAS:GFP* shows strong GFP expression within the lateral plate mesoderm (LPM) along the yolk extension at 26 hpf (white arrows, A,B), which gives rise to the mesothelium at 3-5 dpf (white arrows, C-F). GFP expression is also apparent in subsets of skeletal muscle cells (white arrowheads, A,C,E) and craniofacial muscle cells (yellow arrows, E,F) as well as in cells surrounding the eye (yellow arrowheads, E,F). (G-I) Co-expression of *jam2b^Gt(2A-Gal4)^;UAS:GFP* and *etv2* (red) in the lateral plate mesoderm of the trunk region at 30 hpf. Note that GFP fluorescence and *etv2* expression overlap in the SVF cells (arrowheads), located in the most dorsal region of the GFP expression domain. (J-R) Expression of *jam2b^Gt(2A-Gal4)^;UAS:GFP* in the LPM along the yolk extension and in muscle cells from 30 to 66 hpf. A few GFP-positive cells are present in the PCV and overlap with vascular endothelial marker *kdrl:mCherry* expression (white arrowheads and insets). (S-U) Analysis of *jam2b^Gt(2A-Gal4)^;UAS:GFP* expression in *lyve1b:dsRed* reporter background at 4 dpf, which labels PCV and the thoracic duct. Note GFP-positive cells in the TD (arrowheads). Bright GFP expression in the skeletal muscle fibers is also apparent. PCV, posterior cardinal vein; DA, dorsal aorta; TD, thoracic duct. Lateral views of the trunk region; anterior is to the left. Scale bars: 100 µm.

### *Jam2b*+ cells are derived from the lateral plate mesoderm and incorporate into functional vasculature

To directly visualize the contribution of *jam2b+* cells to the vasculature, we performed time-lapse imaging of *jam2b^Gt(2A-Gal4)^;UAS:GFP*; *Tg(kdrl:mCherry)* embryos starting at 24 hpf. This imaging showed that a subset of GFP-positive cells migrated dorsally away from the LPM region, integrated into the PCV, proliferated, and initiated *kdrl:mCherry* expression as they differentiated into vascular endothelial cells (Fig. 2A-F and Movie 1). In addition, contributions to intersegmental vessels (ISVs) and parachordal lymphangioblasts (PLs) were observed (Movie 2, 3). These data show that mesenchymal cells can contribute to and incorporate into the functional vasculature, even after blood circulation has been initiated, in agreement with our previous observations using the *etv2^Gt(2A-Venus)^* reporter line.^15^

**Figure 2.**
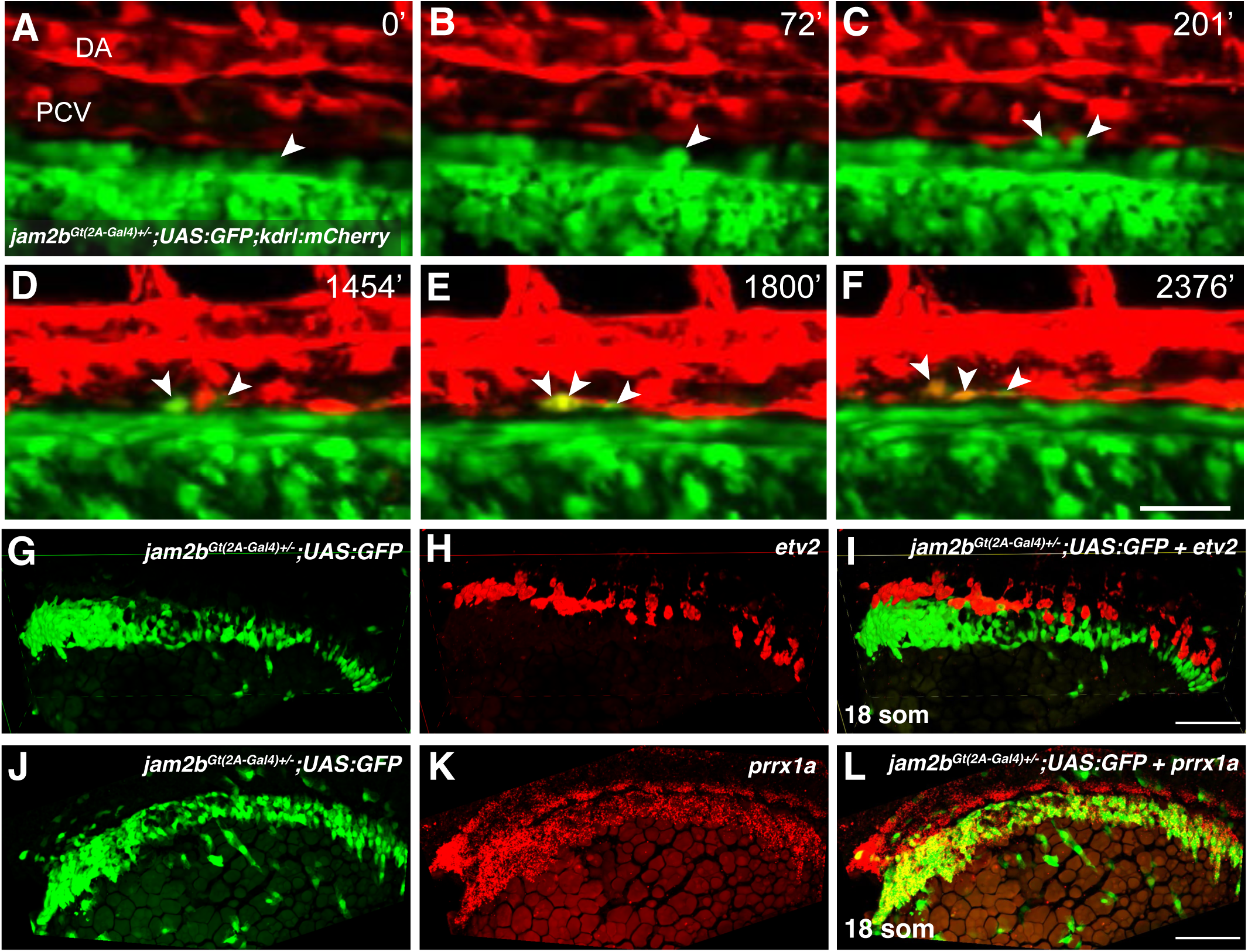
Jam2b-positive cells are derived from the lateral plate mesoderm and contribute to the vasculature. (A-F) Time-lapse images of *jam2b^Gt(2A-Gal4)^;UAS:GFP; Tg(kdrl:mCherry)* embryo starting at 24 hpf show two GFP-positive cells (arrowheads), which migrate dorsally, proliferate, integrate into the PCV and initiate *kdrl:mCherry* expression (see Movie 1). Time is in minutes after 24 hpf. (G-I) HCR against *etv2* in a *jam2b^Gt(2A-Gal4)^;UAS:GFP* embryo at the 18-somite stage showing *jam2b* expression (green) lateral and ventral to *etv2*-positive vascular progenitors (red). (J-L) HCR against *prrx1a* in a *jam2b^Gt(2A-Gal4)^;UAS:GFP* embryo at the 18-somite stage showing colocalization of *prrx1a* mRNA and GFP fluorescence. A dorsolateral view of the trunk region is shown; anterior is to the left. PCV, posterior cardinal vein; DA, dorsal aorta. Scale bar: 50 µm.

In an attempt to identify the origin of SVF cells, we analyzed *jam2b^Gt(2A-Gal4)^;UAS:GFP* expression at the 18-somite stage. Similar to *jam2b* mRNA expression, GFP-positive cells were located bilaterally in the lateral plate mesoderm, mostly lateral and ventral to the emerging *etv2+* vascular progenitors (Fig. 2G-I). There was no apparent overlap between *jam2b^Gt(2A-Gal4)^;UAS:GFP* and *etv2* expression at this stage. This domain largely overlapped with the expression domain of *prrx1a* (Fig. 2J-L), which labels the lateral portion of the LPM.^23^ To analyze the behavior of these cells, we performed time-lapse imaging of *jam2b^Gt(2A-Gal4)^;UAS:GFP* embryos starting at the 18-somite stage. GFP-positive cells remained in the same LPM region from the 18-somite stage to 24-30 hpf (Movie 4). As the embryo continued to undergo convergent extension, the same LPM region became bilaterally located along the yolk extension. Thus, *jam2b*-expressing cells observed at 24-30 hpf along the yolk extension are derived from the ventrolateral portion of the LPM.

### Genetic lineage tracing of *jam2b+* cells

To perform genetic lineage tracing of *jam2b*-expressing cells, we generated *UAS:Cre* and *UAS:CreERT2* DNA constructs. They were coinjected with Tol2 transposase into *jam2b^Gt(2A-Gal4)^; actb2:loxP-BFP-loxP-dsRed; fli1:GFP* embryos, which ubiquitously express floxed BFP under the *actb2* promoter, in combination with the *jam2b^Gt(2A-Gal4)^* knock-in driver and vascular endothelial *fli1:GFP* reporter (Fig. 3A). This approach permanently labels *jam2b*-derived cells, which switch from BFP to dsRed due to Cre-LoxP recombination. 4-hydroxytamoxifen (4-OHT) was added at 7 hpf to UAS:CreERT2-injected embryos to induce recombination. The embryos were then analyzed at 3 dpf by confocal microscopy, revealing switched dsRed-positive cells in different vascular beds. The majority of the labeled cells were present in the SIA, while smaller contributions to the PCV, DA, ISVs, and interconnecting vessels (ICVs) were observed (Suppl. Fig. S5). Due to intense dsRed fluorescence in the mesothelial layer along the yolk and yolk extension we were unable to reliably determine the contribution of *jam2b+* cells to the SIV. Subsequently, we established stable *Tg(UAS:Cre)* and *Tg(UAS:CreERT2)* lines and performed more extensive fate mapping analysis by crossing the *UAS:Cre*-positive adults to *jam2b^Gt(2A-Gal4)^; actb2:loxP-BFP-loxP-dsRed; fli1:GFP*. Confocal microscopy analysis at 3 dpf revealed dsRed expression in the presumptive mesothelial layer surrounding the yolk and yolk extension, a subset of skeletal muscle cells and vasculature (Fig. 3B,E). In addition, both epicardial and myocardial layers within the heart were labeled by dsRed fluorescence (Fig. 3C,D). An embryo negative for *jam2b^Gt(2A-Gal4)^* expression was included for autofluorescence control (Fig. 3H). The majority of labeled vascular endothelial cells were positioned in the SIA and PCV, while a few cells were positioned in the DA, PL, and ISVs (Fig. 3F,G,I-K). Most ISVs that showed dsRed expression were venous, and very little contribution to arterial ISVs was observed (Fig. 3F,G,K). To determine when *jam2b+* cells contribute to vasculature, the *UAS:CreERT2* line was crossed to *jam2b^Gt(2A-Gal4)^; actb2:loxP-BFP-loxP-dsRed; fli1:GFP* adults and 4-OHT was added at different time points to induce Cre-LoxP recombination. 4-OHT addition at 7 hpf mostly labeled the SIA with a smaller fraction of cells present in the SIV, PCV,venous ISVs and PLs (Fig. 3L, Suppl. Fig. S6). When 4-OHT was added at 22 hpf, labeled cells were found mostly in the SIA and, in rare cases, the PCV. Addition of 4-OHT at 7 hpf and subsequent washout at 22 hpf resulted in the labeling of the SIA and a small number of cells in the PCV (Fig. 3L, Suppl. Fig. S6). These results argue that *jam2b+* cells contribute to vasculature both before and after 22 hpf. However, contribution after 22 hpf was largely limited to the intestinal vasculature, which supports our findings that SVF cells contribute substantially to the SIA and SIV.

**Figure 3.**
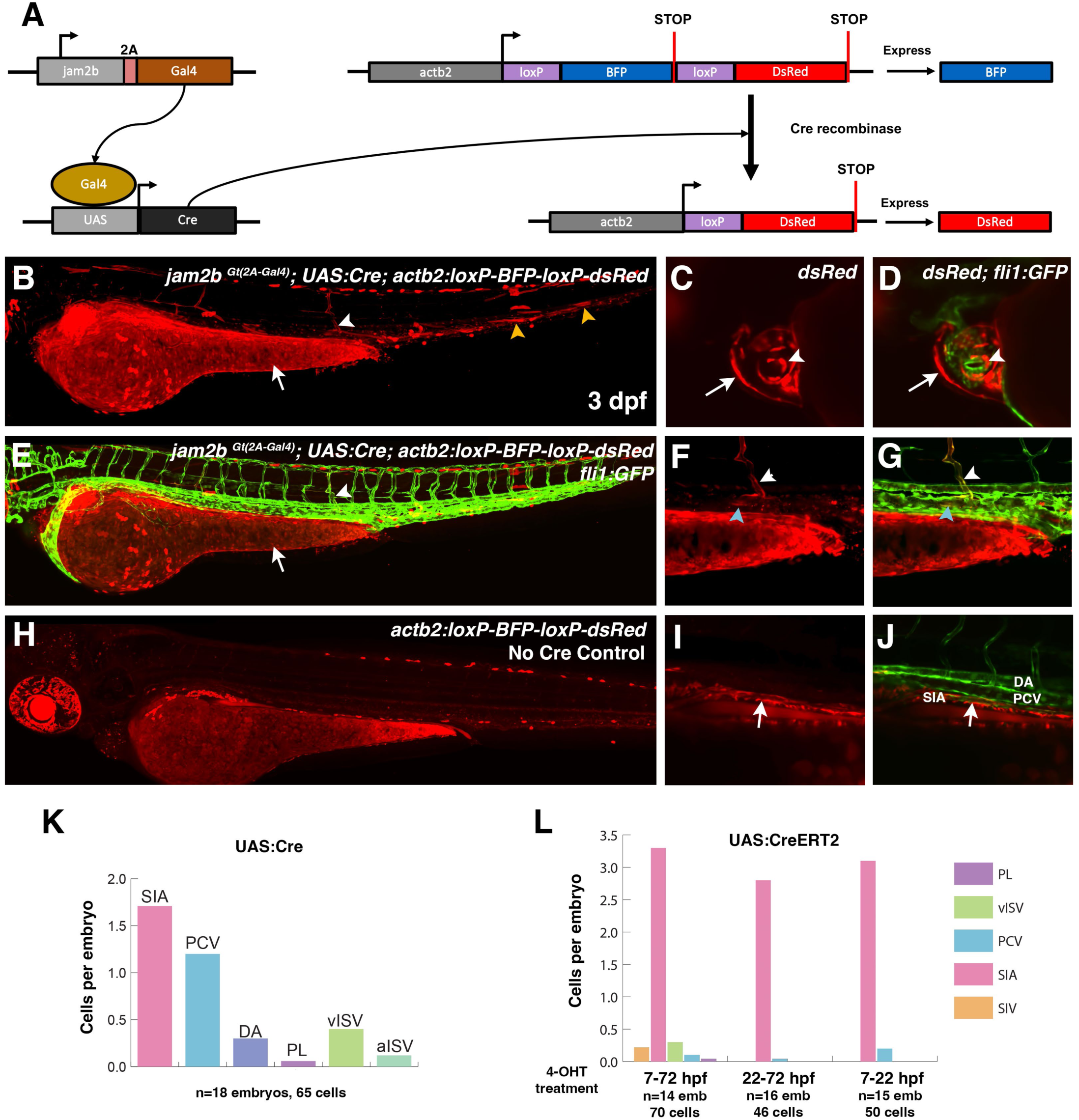
Genetic lineage tracing analysis of *jam2b+* cell contribution to the vasculature. (A) Schematic diagram of recombination in *jam2b^Gt(2A-Gal4);^ UAS:Cre; actb2-loxP-BFP-loxP-dsRed* embryos. Ubiquitous BFP expression switches to dsRed in Cre-positive cells. (B-J) Lineage tracing analysis in embryos at 3 dpf obtained by crossing *jam2b^Gt(2A-Gal4)^*and *UAS:Cre; actb2:loxP-BFP-loxP-dsRed; fli1:GFP* parents. Red cells, indicating the genetic switch, are observed in the mesothelial layer surrounding the yolk (arrow, B,E), skeletal muscle (yellow arrowheads, B), and the epicardial (arrows, C,D) and myocardial (arrowheads, C,D) heart layers. Switched cells in the vasculature are observed in the venous ISVs (white arrowheads, B,E,F,G), the PCV (blue arrowheads, F,G) and SIA (arrow, I,J). A control embryo negative for *jam2b^Gt(2A-Gal4)^* is shown in (H) to indicate autofluorescence. (F,G) show higher magnification views of (B,E). (K) Quantification of endothelial cell labeling in different vascular beds in *jam2b^Gt(2A-Gal4)^*; *UAS:Cre; actb2-loxP-BFP-loxP-dsRed* embryos at 3 dpf. (L) Quantification of endothelial cell labeling in different vascular beds in *jam2b^Gt(2A-Gal4)^*; *UAS:CreERT2; actb2-loxP-BFP-loxP-dsRed* embryos at 3 dpf. 4-OHT was added starting at 7 hpf or 22 hpf. In a third group of embryos, 4-OHT was added at 7 hpf and washed out at 22 hpf. Note that cell contribution to the SIV could not be analyzed reliably due to very strong red fluorescence in the mesothelial layer surrounding the yolk.

Surprisingly, the number of intestinal vascular cells labeled by recombination driven by the stable *jam2b^Gt(2A-Gal4)^; UAS:Cre* or *UAS:CreERT2* lines was rather small (1.5-3 SIA cells per embryo), which was lower than that observed after transient *UAS:Cre* or *CreERT2* expression (average of 6 or 4 SIA cells per embryo). UAS lines are known to undergo silencing, resulting in mosaic expression.^24^ Indeed, ISH analysis of *Cre* expression in *jam2b^Gt(2A-Gal4^; UAS:Cre* embryos revealed a highly variable, mosaic *Cre* expression pattern (Suppl. Fig. S7). This suggests that the true level of contribution of *jam2b-*positive cells to vascular beds is higher than observed in our fate mapping experiments.

### Intestinal vasculature forms largely by new vasculogenesis

Previous studies have argued that the intestinal vasculature, including the SIA and SIV, is derived from the PCV.^8-10^ In contrast, our current data suggest that *jam2b-*positive SVF cells make a significant contribution to the intestinal vasculature. However, the number of intestinal vascular cells labeled by *jam2b^Gt(2A-Gal4)^; UAS:Cre* or *UAS:CreERT2* recombination was rather small, likely due to mosaic *UAS:Cre* and *UAS:CreERT2* expression. To clarify the contribution of new vasculogenesis to the intestinal vasculature, we photoconverted all vascular endothelial cells in the *etv2:Kaede* transgenic line at 24 hpf, which expresses green-to-red photoconvertible Kaede in all vascular endothelial cells,^25^ and then analyzed the percentage of red-and green-labeled cells at 4 dpf in the intestinal vasculature over the yolk extension region, where SVF cells are observed. Notably, 78.5% of cells in the SIA and 80.5% of cells in the SIV showed only green expression of newly synthesized Kaede, and had no red fluorescence from photoconverted protein, arguing that they originated after photoconversion at 24 hpf (Fig. 4). In a control experiment, photoconversion at 3 dpf resulted in red labeling of all intestinal vasculature at 6 dpf (Suppl. Fig. S8), suggesting that little to no new vasculogenesis occurs during these stages and demonstrating that photoconverted Kaede protein is stable for at least 3 days after photoconversion. These data suggest that most intestinal progenitors in the mid and posterior trunk regions form by new vasculogenesis from SVF cells.

**Figure 4.**
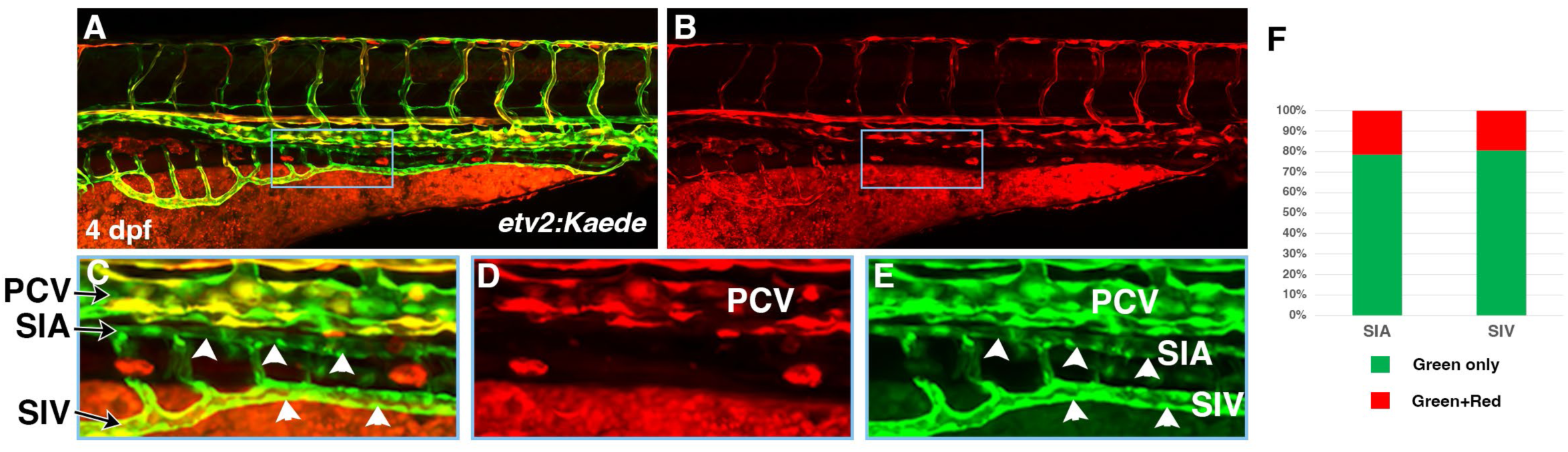
Intestinal vasculature in the middle and posterior trunk region forms largely by new vasculogenesis. (A,B) *etv2:Kaede* embryos were photoconverted from green to red at 24 hpf and analyzed at 4 dpf. Merged and red-only channels are shown. (C-E) Higher magnification images of the boxed area in (A,B). Merged, red and green channels are shown. Note that SIA and SIV are composed mostly of green-only cells (white arrowheads), indicating that they originated after the photoconversion. (F) Quantification of green-only (new) and green+red (pre-existing) cells in the SIA and SIV at 4 dpf. The analysis focused on the middle and posterior trunk region over the yolk extension area.

### Transcriptomic analysis of SVF cells

We have previously performed single-cell RNA sequencing (scRNA-seq) of the entire trunk region of zebrafish embryos in order to identify the transcriptional profile of SVF cells.^15,16^ While this analysis identified a set of SVF marker genes that included *etv2, tal1, lmo2* and *jam2b,* only a limited number of marker genes were identified with this approach, because SVF cells comprise only a small portion of cells in the entire trunk. Therefore, we used *jam2b* and *etv2* reporter lines to perform additional transcriptomic analysis in order to characterize the profiles of SVF cells and their transition to vascular endothelial cells in greater detail. mCherry and Venus-positive cells were FACS-sorted from *jam2b^Gt(2A-Gal4)^;UAS:mCherry-NTR; etv2^Gt(2A-Venus)^* embryos at 36 hpf and subjected to scRNA-seq using the Chromium platform (10x Genomics). Bioinformatic analysis using Seurat 4.0 identified a total of 37 cell clusters, including the SVF cell cluster, five venous, three arterial clusters, and a capillary endothelial cluster (Fig. 5A-C, Suppl. Table 1). The SVF cell cluster (cluster #15) was identified based on top marker genes, which included *npas4l, tal1, lmo2,* and *etv2* (Fig. 5D-F, Suppl. Table 1). Additional top SVF marker genes included *gfra3, sox7, si:dkey-52l18.4, egfl7* and *tmem88a* (Suppl. Table 1). A subset of cells in cluster 15 was also positive for *jam2b* expression (Fig. 5D,E). It is likely that the cluster 15 includes SVF cells at different stages of differentiation toward vascular endothelium, while *jam2b* expression is low or absent in differentiated vascular endothelial cells, which explains why only a subset of cells in cluster 15 show overlapping *etv2* and *jam2b* expression. The key SVF marker genes, including *etv2, tal1, lmo2, sox7, si:dkey-52l18.4* and *egfl7* were also identified in our earlier study,^15^ arguing that the current approach has successfully identified the transcriptional signature of SVF cells.

**Figure 5.**
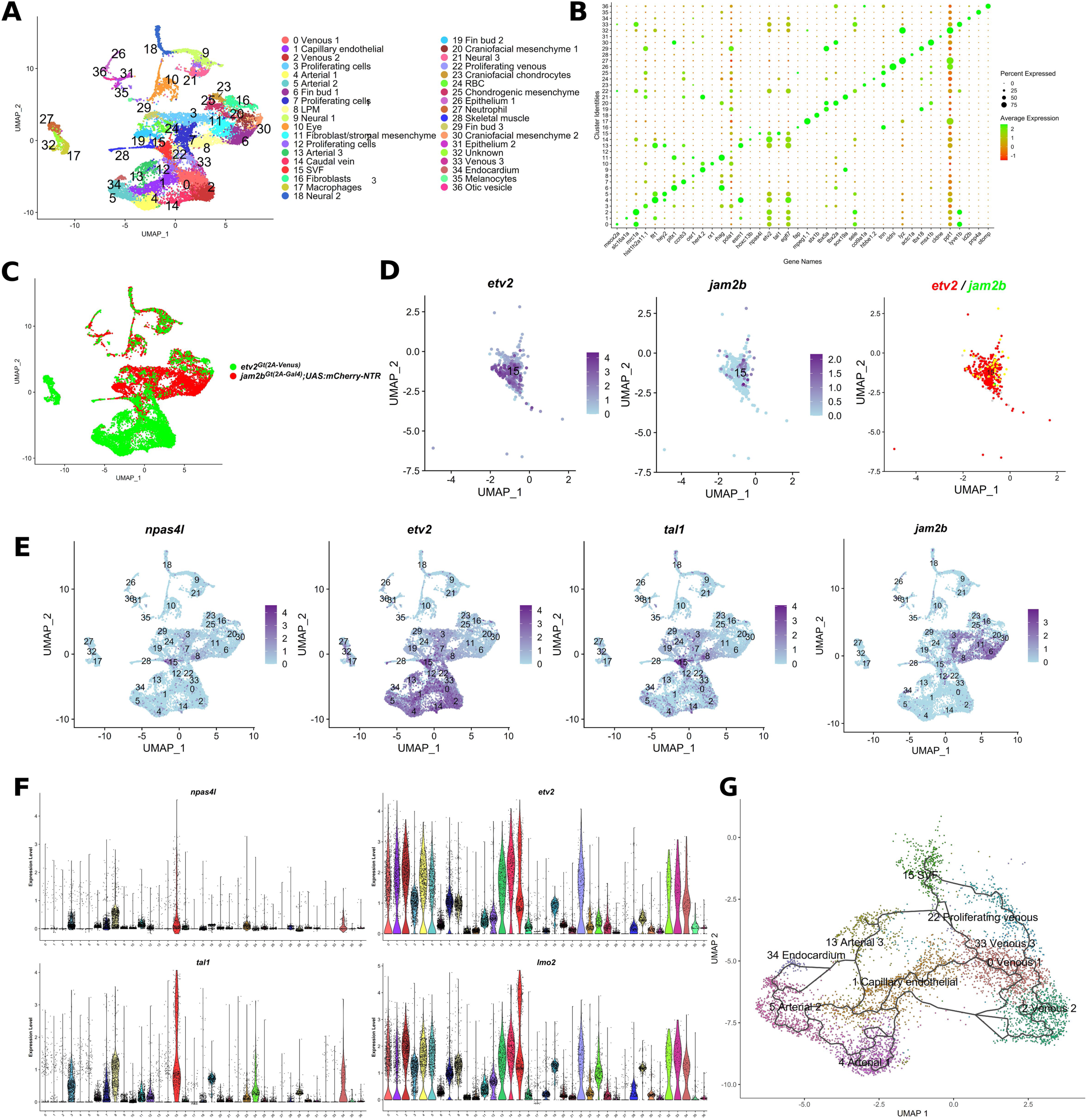
scRNA-seq analysis of cells from *jam2b^Gt(2A-Gal4)^;UAS:mCherry-NTR; etv2^Gt(2A-Venus)^* embryos at 36 hpf. (A) UMAP plot showing 37 cell clusters identified by Seurat analysis. LPM, lateral plate mesoderm; SVF, secondary vascular field; RBC, red blood cells. (B) Dot plot of selected marker gene expression levels in each cell cluster. (C) 2D UMAP plot showing the distribution of Venus and mCherr*y* positive cells. Note that Venus-positive cells mostly correspond to different vascular endothelial lineages, while mCherry-positive cells label different subsets of LPM. (D) Feature plots of cluster 15 showing expression of *etv2, jam2b* and coexpression of *etv2* and *jam2b* (yellow). Note that only a subset of cells coexpress *etv2* and *jam2b* (E) Feature plots showing the expression of SVF cell markers, enriched in the cluster 15. (F) Violin plots show SVF cell marker expression in different clusters. (G) Pseudotime trajectory of vascular endothelial and SVF clusters generated using Monocle 3. Note that SVF cells (cluster 15) can transition into both venous and arterial cell types.

To identify transitional events during SVF cell differentiation into vascular endothelium, we performed cell lineage trajectory analysis using Monocle 3.^26^ The cell lineage trajectory showed that SVF cells can directly transition into both venous and arterial clusters (Fig. 5G). This differentiation trajectory is consistent with experimental observations that SVF cells contribute to both venous (PCV, SIV) and arterial (SIA) vasculature. Unfortunately, it is currently not possible to distinguish SIA or SIV-specific cells on the lineage trajectory from other types of arterial or venous cells due to the absence of specific markers for these cells.

### *Jam2b* mutants undergo normal vascular development

To analyze the functional role of Jam2b in vascular development, we generated *jam2b* mutants using CRISPR/Cas9-mediated genome editing. Two different guide RNAs were used to delete the *jam2b* promoter region, and a stable mutant line was established (Suppl. Fig. S9A). As indicated by ISH analysis, *jam2b-/-* embryos showed no detectable *jam2b* expression in the LPM region, suggesting that the mutant may be a null or close to a null (Suppl. Fig. S9C,D). However, both zygotic and maternal-zygotic *jam2b-/-* embryos appeared morphologically normal and did not show any apparent vascular defects based on *kdrl:GFP* expression analysis (data not shown). *Jam2b* homozygous mutants were viable as adults and did not exhibit any apparent defects. *Etv2* expression in SVF cells was also not affected in maternal-zygotic *jam2b* mutant embryos (Suppl. Fig. S10A-C). We have previously demonstrated that SVF cells contribute to vascular recovery after endothelial cell ablation in *etv2^Gt(2A-Gal4)^; UAS:mCherry-NTR* embryos.^15^ We then tested whether vascular recovery was affected in *jam2b*-deficient embryos. Metronidazole (MTZ) was added at approximately 10 hpf stage and then washed out at 45 hpf stage in zygotic *jam2b-/-* and sibling control *jam2b+/-* embryos in an *etv2^Gt(2A-Gal4)^; UAS:mCherry-NTR; kdrl:GFP* background. As we previously demonstrated, such treatment results in a complete ablation of all vascular endothelial cells except for SVF cells, which emerge later than other endothelial cells.^15^ Afterwards, recovery was analyzed at 72 hpf, when the partially regenerated vascular cords were apparent; as our previous study suggested, these cords were derived from SVF cells.^15^ However, no difference in vascular recovery was observed between *jam2b-/-* and *jam2b+/-* embryos (Suppl. Fig. S10D,E).

### *jam2a; jam2b* mutants display defects in SVF cell formation and intestinal and venous vasculature

We then explored the possibility that *jam2b* may function redundantly with the related homolog *jam2a.* While *jam2a* is broadly expressed in the somitic muscle,^20,21,27^ our scRNA-seq analysis indicated that it also showed significant expression in the SVF cell population, although it was not enriched among marker genes.^15^ ISH analysis demonstrated that *jam2a* expression is concentrated in perivascular regions along the axial vessels and ISVs and overlaps with *etv2* expression in SVF cells (Suppl. Fig. S11). We used CRISPR / Cas9 mutagenesis to remove the entire coding sequence of *jam2a* (Suppl. Fig. S9B,E,F). To expedite the generation of double mutant embryos, *jam2a* mutant generation was performed in a *jam2b-/-* mutant background. A stable *jam2a;jam2b* mutant line was generated and used for further experiments. Zygotic *jam2a-/-;* maternal-zygotic *jam2b-/-* mutant embryos, obtained by the in-cross of *jam2a+/-;jam2b-/-* adults did not show any apparent defects in vascular development or SVF cell formation (data not shown). Subsequently, maternal-zygotic (MZ) mutant embryos were obtained and analyzed for SVF cell formation and vascular defects. Interestingly, MZ *jam2a-/-;jam2b-/-* embryos showed a significant reduction in the SVF cell number when compared to control double heterozygous embryos in the same background. The number of *etv2, tal1* and *npas4l-*positive SVF cells was greatly reduced in MZ *jam2a;jam2b* mutant embryos, while no other morphological defects were apparent (Fig. 6). Overall vascular patterning appeared largely unaffected in MZ *jam2a;jam2b* mutant embryos based on *kdrl:GFP* fluorescence analysis at 3 dpf (Fig. 7A,B). However, MZ *jam2a;*jam2b mutant embryos showed a greater incidence of incompletely extended ISVs (Fig. 7A-C). Venous ISVs, derived from the PCV, accounted for 97% (33 out of 34) of such truncated ISVs, while only 3% were arterial. This suggested that the number of venous cells could be reduced, possibly due to reduced contribution of SVF cells to the PCV. Indeed, the number of ECs in the floor of the PCV was slightly but significantly reduced in MZ *jam2a; jam2b* mutant embryos (Fig. 7D-F). However, the most pronounced defects were observed in the formation of the intestinal vasculature, particularly the SIA, which was underdeveloped and showed gaps or missing segments in MZ *jam2a;jam2b* mutants (Fig. 7G-I). In addition, a significant fraction of MZ *jam2a;jam2b* mutants showed gaps in the SIV (Fig. 7G-I). To test the contribution of new vasculogenesis to the PCV and intestinal vasculature, we photoconverted *fli1:Kaede* at 24 hpf in MZ *jam2a; jam2b* mutant and control embryos, and then analyzed the presence of new endothelial cells at 3 dpf. MZ *jam2a; jam2b* mutants showed greatly increased fraction of red *fli1:Kaede+* cells in the SIA compared to control embryos, while the number of green-only cells in the PCV, that originate by new vasculogenesis, was significantly reduced (Suppl. Fig. S12). These results argue that the formation of new vascular progenitors and their contribution to venous and intestinal vasculature is greatly reduced in MZ *jam2a, jam2b* mutants.

**Figure 6.**
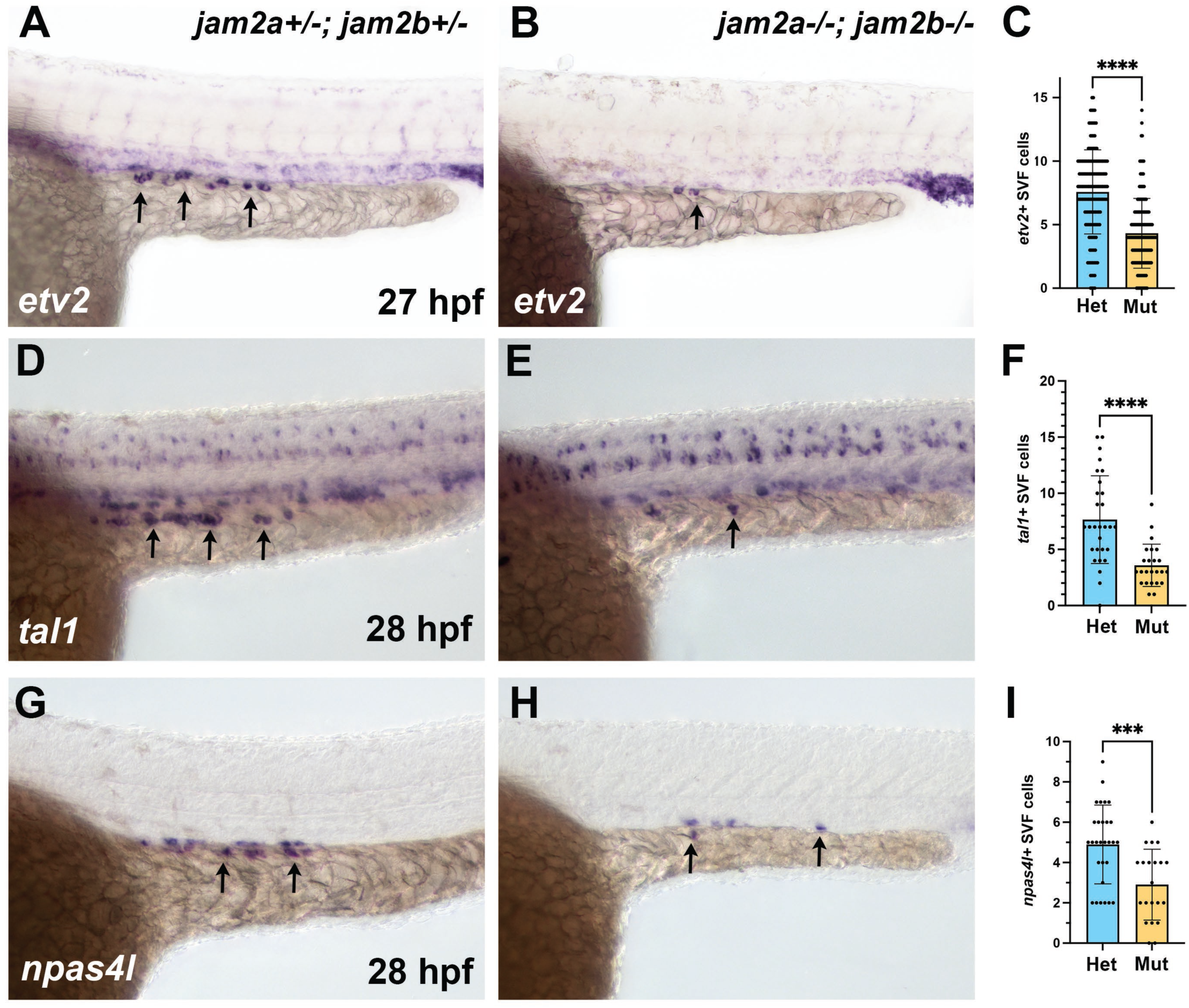
SVF cell number is reduced in maternal-zygotic (MZ) *jam2a; jam2b* mutant embryos. (A-C) In situ hybridization analysis (ISH) for *etv2* expression in SVF cells (arrows, A,B) in MZ *jam2a-/-; jam2b-/-* embryos and control *jam2a+/-; jam2b+/-* embryos in *kdrl:GFP* background, obtained by crossing double homozygous *jam2a-/-; jam2b-/-; kdrl:GFP* adults to wild-type *kdrl:GFP*. (D-I) ISH analysis for *tal1* and *npas4l* expression in SVF cells (arrows) in MZ *jam2a-/-; jam2b-/-* embryos and control embryos in *fli1:GFP* background. Controls were obtained by mating *jam2a+/-; jam2b-/-* embryos to wild-type *fli1:GFP,* resulting in a mixture of *jam2a+/+; jam2b+/-* and *jam2a+/-; jam2b+/-* genotypes. ***p<0.001, ****p<0.0001, two-tailed t-test; error bars indicate ± s.d. Data are combined from four (C) or two (F,I) independent replicate experiments. Het labels in (C,F,I) refer to double heterozygous in (C) and a mixture of *jam2a+/+; jam2b+/-* and *jam2a+/-; jam2b+/-* genotypes in (F,I), while the Mut label refers to *jam2a-/-; jam2b-/-* MZ mutant embryos.

**Figure 7.**
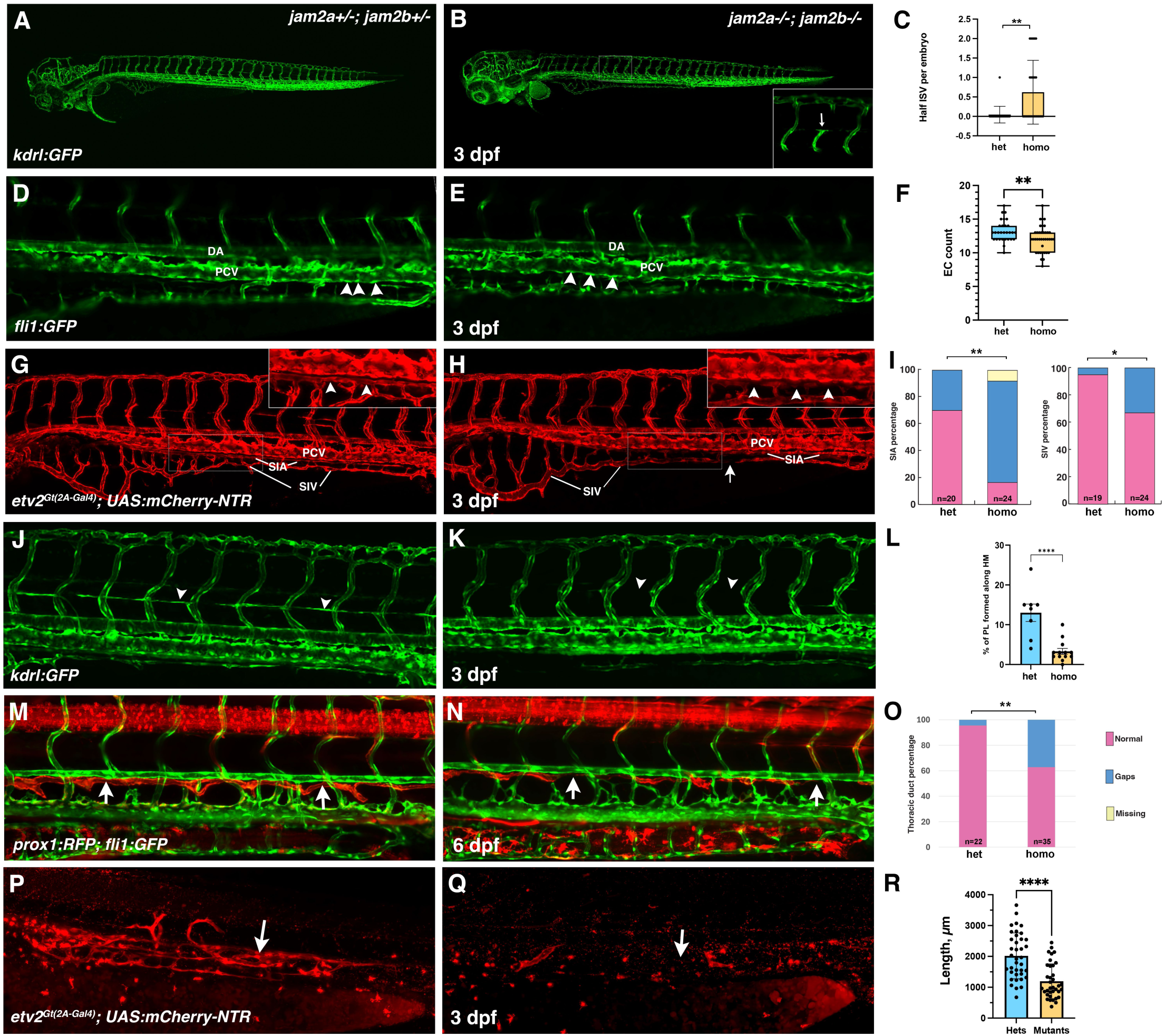
Analysis of vascular defects in MZ *jam2a; jam2b* mutant embryos. (A-C) Intersegmental vessel extension analysis in MZ *jam2a-/-; jam2b-/-* and *jam2a+/-; jam2b+/-* embryos in *kdrl:GFP* background at 3 dpf. Note that some ISVs are truncated in MZ *jam2a; jam2b* mutant embryos (arrow, B; the inset shows a higher magnification view of the boxed trunk region). (D-F) Endothelial cell number in the PCV is reduced in MZ *jam2a; jam2b* mutant embryos compared to *jam2a+/-; jam2b+/-* controls at 3 dpf. Endothelial cells were counted in the floor of the PCV (arrowheads) in the selected area of the trunk region based on *fli1:GFP* expression. (G-I) SIA and SIV are fragmented and show an increased incidence of gaps in MZ *jam2a;jam2b* mutant embryos compared to heterozygous controls at 3 dpf. Analysis was performed in the background of *etv2^Gt(2A-Gal4)^; UAS:mCherry-NTR* transgene, which shows expression in all vasculature. Note the underdeveloped SIA (arrowheads) and gaps in SIV (arrow) in *jam2a;jam2b* mutants. Higher magnification views are shown in the insets. Diagrams in (I) show the percentage of embryos with gaps in SIA (left) and SIV (right). (J-L) Analysis of parachordal lymphangioblasts (PLs) in *kdrl:GFP* transgenic embryos at 3 dpf. Note that PLs at the horizontal myoseptum (arrowheads) are largely absent in MZ *jam2a; jam2b* mutants. Quantification in (L) shows the percentage of the horizontal myoseptum covered by PLs. (M-O) Thoracic duct analysis in *prox1:RFP; fli1:GFP* transgenic embryos at 6 dpf. Note the gaps in the thoracic duct (arrows) observed in MZ *jam2a;jam2b* mutant embryos. (O) Quantification of the percentage of embryos with normal or fragmented thoracic duct. (P-R) Vascular recovery analysis after endothelial cell ablation. *jam2a; jam2b* mutant and control heterozygous embryos in *etv2 ^Gt(2A-Gal4)^; UAS:NTR-mCherry* background were treated with MTZ starting at 6 hpf to ablate ECs and subsequently washed out at 45 hpf. A partial vascular recovery is observed in the control embryos, while it is greatly reduced in MZ *jam2a;jam2b* mutants. (R) Quantification of the length of the recovered vasculature. Data in (C,F,L,R) were analyzed using two tailed t-tests, while data in (I,O) were analyzed using Fisher’s exact tests. *p<0.05, **p<0.01, ****p<0.0001. DA, dorsal aorta; PCV, posterior cardinal vein; SIA, supraintestinal artery; SIV, subintestinal vein.

Because our data suggested that *jam2b+* cells may contribute to lymphatics, we analyzed the formation of the parachordal lymphangioblasts and the thoracic duct using *kdrl:GFP* and *prox1:RFP; fli1:GFP* reporter embryos, respectively. The number of parachordal lymphangioblasts was greatly reduced in MZ *jam2a; jam2b* mutants at 3 dpf compared to control *jam2a+/-;jam2b+/-* embryos (Fig. 7J-L). Furthermore, approximately 37% of MZ *jam2a;jam2b* mutants had gaps in the thoracic duct compared to 5% of *jam2a+/-;jam2b+/-* embryos (Fig. 7M-O). These results argue that *jam2a* and *jam2b* functions are required for the formation of the intestinal vasculature and lymphatics.

As described above, our previous work showed that SVF cells participate in vascular recovery after endothelial cell ablation. We therefore tested whether vascular recovery after endothelial cell ablation was affected in MZ *jam2a; jam2b* mutant embryos in an *etv2^Gt(2A-Gal4)^; UAS:mCherry-NTR* background. In order to ablate all vascular endothelial cells, MTZ was added at 6 hpf and then washed out at approximately 45 hpf. Subsequently, embryos were analyzed at 72 hpf by confocal imaging. Substantial recovery of the PCV and intestinal vasculature was observed in the control *jam2a; jam2b* double heterozygous embryos, whereas recovery was greatly reduced in MZ *jam2a; jam2b* mutant embryos (Fig. 7P-R). Altogether, these results argue that the loss of SVF cells in MZ *jam2a; jam2b* mutants leads to defects in venous, intestinal and lymphatic vessel development, and is also associated with reduced vascular recovery after EC ablation.

### *Hand2* functions upstream of *jam2b* in SVF cell formation

Recent work has shown that the bHLH transcription factor *hand2,* previously implicated in cardiac development, is also expressed in the lateral plate mesoderm along the yolk extension and labels mesothelial and pericardial progenitors, similar to *jam2b.*^28,29^. Intriguingly, *hand2-2A-Cre-*based lineage tracing showed contributions to the venous and intestinal vasculature,^30^ similar to our results obtained using *jam2b^Gt(2A-Gal4)^; UAS:Cre*. To determine whether both genes are co-expressed in the same cells, we performed fluorescent hybridization chain reaction (HCR) analysis at 24 hpf. Indeed, expression of *hand2* and *jam2b* largely overlapped in the LPM region along the yolk extension (Fig. 8A-F). To test whether *hand2* may function upstream of *jam2b,* we analyzed *jam2b* expression in the previously generated *hand2* mutants by in situ hybridization. Indeed, *jam2b* expression was absent in *hand2* mutants from most of the LPM, including the SVF-forming region along the yolk extension (Fig. 8G-J). Only the anterior domain of *jam2b* expression adjacent to the yolk was not affected in *hand2* mutants (Fig. 8G-J, white arrowheads). We then analyzed whether *hand2* function is required for *etv2* expression in SVF cells. Indeed, *hand2* mutants showed a complete absence of SVF cells at 30 hpf (Fig. 8K,L), which correlated with the absence of SIA and SIV at 72 hpf (Suppl. Fig. S13). In addition, *hand2* mutants showed reduced *etv2* expression in the PCV at 30 hpf and partially fused DA and PCV at 72 hpf, consistent with the previously demonstrated requirement for *hand2* in the formation of venous endothelium.^31^

**Figure 8.**
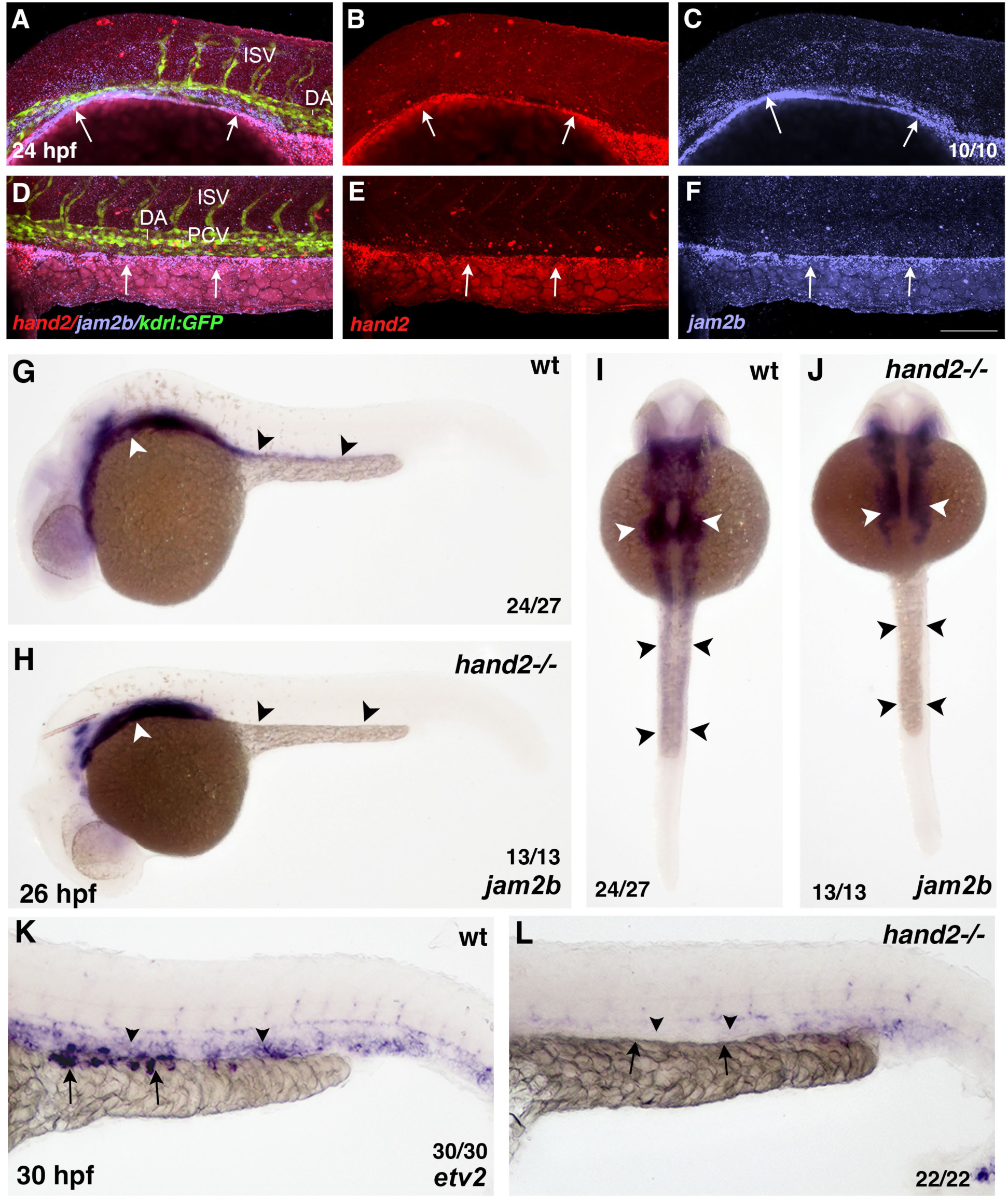
*hand2* is required for *jam2b* expression in the lateral plate mesoderm and SVF cells. (A-F) HCR analysis for *hand2* (red) and *jam2b* (magenta) expression in *kdrl:GFP* embryos at 24 hpf. Anterior (A-C) and posterior (D-F) regions of the trunk are shown. Note the overlapping expression in the LPM along the yolk and yolk extension (arrows), which does not overlap with *kdrl:GFP* expression. (G-J) Whole-mount in situ hybridization analysis for *jam2b* expression in *hand2* mutant and wt control embryos at 26 hpf. Lateral (G,H) and dorsal (I,J) views. Note that *jam2b* expression in *hand2* mutants is absent from the posterior domain of the LPM along the yolk extension (black arrowheads), while the anterior portion of the LPM (white arrowheads) is only mildly affected. (K,L) Whole-mount in situ hybridization for *etv2* expression in *hand2* mutant and control wild-type embryos at 30 hpf. Note that *etv2* expression in SVF cells (arrows) is absent in *hand2* mutants. In addition, its expression in the PCV (arrowheads) is greatly reduced or absent. ISV, intersegmental vessel; DA, dorsal aorta; PCV, posterior cardinal vein. Scale bars: 100 μm.

We then aimed to test whether *jam2b* overexpression was sufficient to restore SVF cell formation in the absence of *hand2* function. First, we tested whether *jam2b* overexpression was sufficient to induce ectopic SVF cell formation in wild-type embryos. Indeed, in situ hybridization analysis for *etv2* expression showed that SVF cell number was significantly increased in wild-type embryos injected with *jam2b* mRNA (Suppl. Fig. S14A-C). However, *jam2b* overexpression in *hand2* mutants failed to rescue any SVF cell formation (Suppl. Fig. S14D). These results show that *hand2* function is required for both *jam2b* expression in the LPM region and SVF cell specification.

## DISCUSSION

Our previous work has demonstrated that new *etv2+* progenitors emerge from the trunk region of the LPM along the yolk extension and contribute to the intestinal and axial vasculature.^15^ Here, we implicate *jam2a* and *jam2b* in the emergence of SVF cells and their contribution to vasculature. Our results show that *jam2b* is expressed across a broad region of the LPM, including the site where SVF cells emerge. Lineage tracing studies revealed that *jam2b-* expressing cells contribute substantially to the intestinal vasculature and to a lesser extent to the PCV and venous ISVs (Fig. 9). MZ j*am2a;jam2b* mutants show a reduced number of SVF cells and display defects in the intestinal and venous vessel development. We further show that the *hand2* transcription factor functions upstream of *jam2b* in the specification of SVF cells within the trunk region of the LPM.

**Figure 9.**
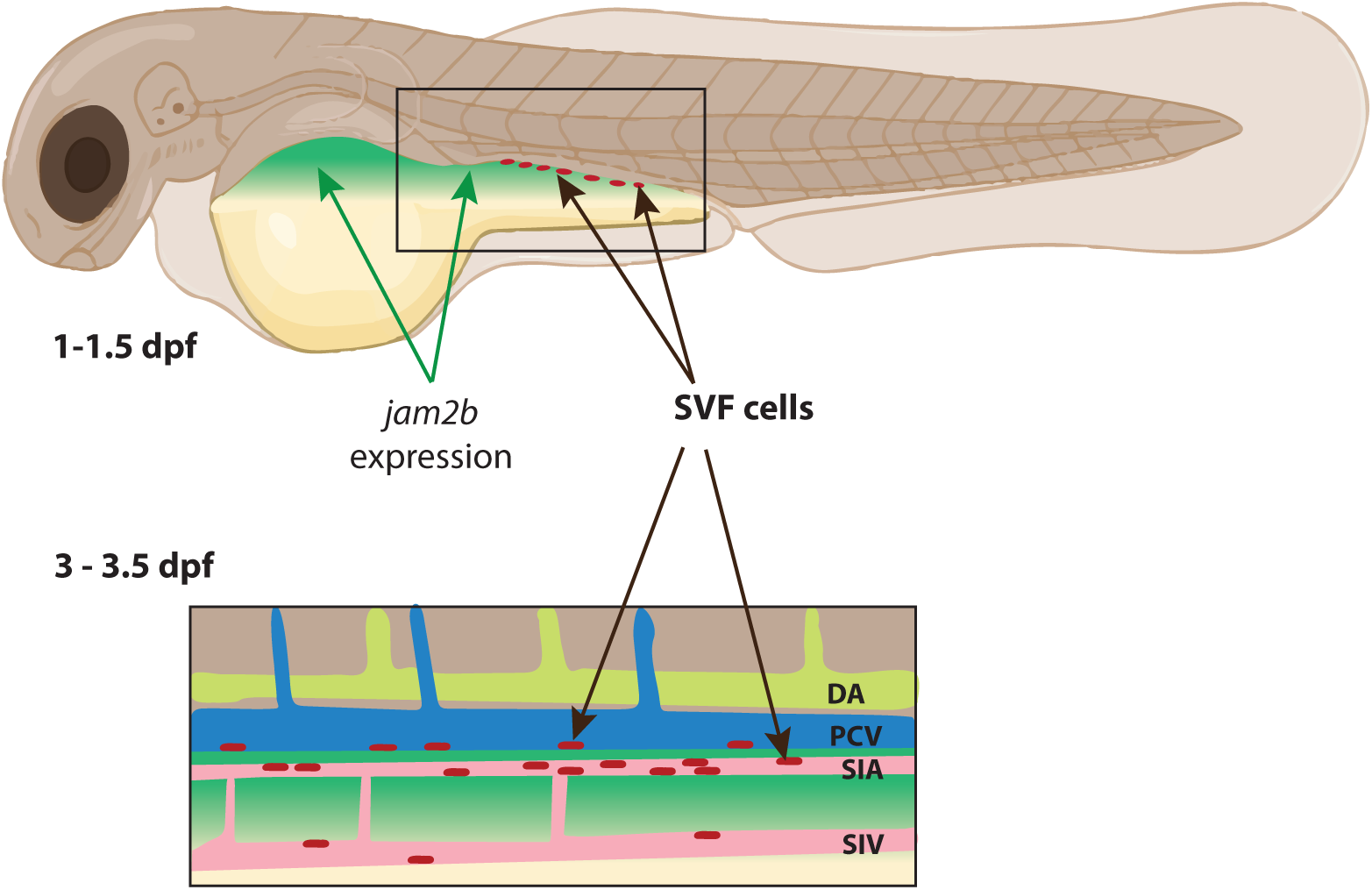
A diagram illustrating the emergence of SVF cells from *jam2b*-positive lateral plate mesoderm. SVF cells contribute to the intestinal vasculature, including the supraintestinal artery (SIA) and subintestinal vein (SIV), as well as the posterior cardinal vein (PCV).

Previous studies have argued that internal organ-specific vessels, including the intestinal vessels, are derived from the PCV.^8-10^ According to a recent model, individual cells migrate and assemble into the two vessels, the SIA and SIV, which can further branch and give rise to ICVs.^8-10^ Our recent study challenged this model and showed that SVF cells can contribute to intestinal vasculature.^15^ However, it has not been clear whether SVF cells make only a minor contribution to the intestinal vasculature. Our current results suggest that SVF cells may be the main source of the intestinal vasculature, while the PCV makes only a minor contribution. *Etv2:Kaede* photoconversion showed that most cells in the intestinal vasculature emerge after the 24 hpf stage and, therefore, they were not present in the PCV at the time of the photoconversion. Although *jam2b^Gt(2A-Gal4);^ UAS:Cre*-based lineage tracing resulted in switched dsRed expression in only a fraction of the PCV and intestinal vessels, this is likely due to the mosaic expression of Cre, as evident from our ISH analysis for *Cre* expression. Based on our earlier time-lapse lineage tracing experiments, it appears that some SVF cells migrate directly to the SIA and SIV, while others integrate into the PCV first, and only afterward exit the PCV and migrate to the intestinal vasculature.^15^ Thus, our model can be reconciled with the previous observations if some SVF cells first integrate into the PCV and subsequently contribute to the intestinal vasculature. It is also possible that the earlier studies focused largely on the anterior portion of the intestinal vasculature, which forms the plexus-like structure and is largely derived from the PCV. Our study focused on the mid-trunk region along the yolk extension, where SVF cells are positioned, which is posterior to the intestinal plexus analyzed in the previous studies.

*jam2b* expression within the LPM is observed during somitogenesis. It is evident that this region gives rise to diverse tissues, including the mesothelium, pectoral fins, pericardium, and muscle. A similar contribution has also been demonstrated for *hand2*-positive LPM cells,^29^ which largely overlap with *jam2b* expression. Only a small fraction of *jam2b*-positive cells contributes to the vascular endothelium. The cells fated to become endothelium are located at the edge of the medial portion of the *jam2b* expression domain in the trunk region, closest to the midline (Fig. 9). It is tempting to speculate that all *jam2b-*positive cells are multipotent initially, yet they experience different signals based on their relative position. The cells closest to the midline in the trunk region are induced to differentiate into vascular endothelium. The inductive signal is currently unknown and will require further investigation. Interestingly, *vegfaa* is expressed in the region where intestinal vessels form.^8^ Furthermore, *vegfaa* mutants show a reduced number of SVF cells, which correlates with the defects in the intestinal vessel formation,^8,15^ suggesting that *vegfaa* could be a candidate signal.

During embryonic vasculogenesis arterial and venous progenitors originate from the LPM during somitogenesis.^11-14^ While the arterial progenitors originate close to the midline, venous progenitors emerge from the more lateral portion of the LPM.^25^ As we show here, early venous progenitors emerge adjacent to *jam2b*-positive LPM cells, which are positioned further lateral to the venous progenitors. While the expression domains of *jam2b* and *etv2* overlap in later-forming SVF cells, they do not appear to overlap in early endothelial progenitors, which was also corroborated in our scRNA-seq data.^15,23^ We also did not observe *jam2b^Gt(2A-Gal4)^; UAS:GFP* expression in the PCV before 24 hpf, suggesting that *jam2b* may not be involved in early vasculogenesis. It appears that *jam2b+* LPM cells maintain vasculogenic potential beyond 24 hpf.

The mechanism by which *jam2a* and *jam2b* regulate SVF cell specification is unclear. A previous study has demonstrated that antibodies against murine Jam-B (Jam2) inhibited angiogenesis in vitro and interfered with the VEGFR2 signaling pathway.^32^ Our previous data indicate that VEGF signaling is required to upregulate *etv2* expression during early vasculogenesis, and it is also required for SVF cell emergence.^15,33^ It is possible that Jam2 promotes VEGF signaling, a possibility that will require investigation in future studies.

Previous studies have demonstrated that *hand2* function is required for PCV development. *npas4l* and *etv2* expression were reduced in the venous progenitors, arguing that *hand2* functions upstream of these transcription factors.^31,34^ We show here that SVF cells are also absent in *hand2* mutants. It is possible that *hand2* directly regulates *npas4l* and/or *etv2* expression. Alternatively, *hand2* may function through *jam2b* and indirectly be involved in SVF specification through VEGF or a different pathway. Because *jam2b* overexpression in *hand2* mutants fails to rescue *etv2* expression in SVF cells, this finding suggests that *jam2b* function is not sufficient for SVF cell specification in the absence of *hand2*.

In summary, this study demonstrates that the *jam2b-*positive LPM serves as a major source of internal organ-specific vascular progenitors and identifies the functional requirement for *jam2a* and *jam2b* in promoting *etv2* expression and vasculogenesis. Because analogous late-forming vascular progenitors have also been identified in mammalian embryos,^15^ it is likely that the mechanisms regulating their emergence are also evolutionarily conserved. These results will be important for understanding the molecular mechanisms involved in the formation of different vascular beds and will promote the development of new approaches aimed at vascular regeneration.

## MATERIALS AND METHODS

### Zebrafish lines

The following previously established lines were used in the study: wild-type AB (Zebrafish International Resource Center), *Tg(kdrl:mCherry)^ci5^,*^35^ *Tg(fli1a:GFP)^y1^,*^36^ *hand2^s6^,*^28^ *etv2^Gt(2A-Gal4)ci32^;UAS:mCherry-NTR,*^37^ *etv2^Gt(2A-Venus)^,*^15^ *Tg(actb2:loxP-BFP-loxP-dsRed)^sd27^,*^38^ *Tg(kdrl:GFP)^s843^,*^14^ *TgBAC(prox1a:KalT4-UAS:uncTagRFP)^nim5^* ^39^ (abbreviated as *prox1:RFP*), *Tg(lyve1b:dsRed)^nz101^.*^40^

### Generation of jam2b^Gt(2A-Gal4)^ knock-in line

To generate the *jam2b^Gt(2A-Gal4)ci55^;UAS:GFP* knock-in line (further abbreviated as *jam2b^Gt(2A-Gal4)^*; UAS:GFP), a gBlock Gene Fragment was synthesized by IDT and included the *jam2b* sgRNA site, the P2A sequence, the *Gal4* DNA binding domain and a poly-A tail. This fragment was then A-tailed using dATP and Taq DNA Polymerase (NEB, cat. no. M0273) followed by TA cloning into the TOPO dual vector according to manufacturer’s instructions (ThermoFisher, cat. no. 450640). A mixture of 300 pg *Cas9* mRNA and 50 pg *jam2b-sgRNA-P2A-Gal4* plasmid DNA was injected into Tg(*UAS:GFP*)^41^ embryos at the one-cell stage. Embryos showing any specific *GFP* expression were raised through adulthood and screened to identify a founder which produced progeny with a specific expression pattern.

### Generation of jam2a and jam2b mutant lines

To generate the *jam2b^sfl2^* line, two synthetic gRNAs (Synthego) targeting the *jam2b* promoter region were co-injected with Cas9 protein (NEB, EnGen Spy Cas9 NLS, Cat No. M0646) into zebrafish embryos at the one-cell stage. The following sgRNA sequences were used: GCAATTCTGGAAGAAGTCAA and AATAAGCAGGAGAAGAGCAG. Successful deletion was confirmed using the following primers outside of the deletion site: *jam2b_P3.4.5_F:* GCATCACTCAAAGGCAAATG and *jam2b_P1.2_R:* ACAGTTCCCACCATTTCAGC. A single founder with a 1,071 bp deletion that encompasses the *jam2b* proximal promoter region and a portion of exon 1, was identified and used to establish the mutant line. The genomic sequence flanking the deleted region is shown here: TATAGACCGGGCAGATTAGCCATTG………TCTTCTCCTGCTTATTTACAGTAAG.

To generate the *jam2a^sfl3^* line, two synthetic gRNAs (Synthego) were designed to delete the *jam2a* coding sequence with the following sequences: TCTGTGTGCCTCTAGGTGAG and CAGAGCACAAACCCACGCAA. Both gRNAs were coinjected together with Cas9 protein into one-cell stage zebrafish embryos, homozygous for the *jam2b^sfl2^*mutation. Successful deletion was confirmed using the following primers that flank the deletion site: *jam2a-sg1_2_F:* CAAACACCTGTGACCTTACAAAA and *jam2a-sg1_2_R:* TATCAATGGCAACACAGTGG. A founder carrying a 15,771 bp deletion that includes the entire *jam2a* coding sequence was identified and used to establish the stable line for further studies. The genomic sequence flanking the deleted region is shown here: CCTACAGGTGGTTTGTCAAGCCCAC AAAGGTAGGGAGAACATGCAAACTC

### Generation of UAS:Cre and UAS:CreERT2 lines

To generate *UAS:Cre^sfl4^* and *UAS:CreERT2^sfl5^*lines, the gene fragment containing 5x UAS sequence, *hbae3* 5’and 3’UTR sequences, a zebrafish Kozak sequence,^42^ NLS-Cre, and NLS-CreERT2 codon optimized for zebrafish expression using the iCodon approach,^43^ the SV40 polyA site, and attL4 and attL3 sequences specific to LR recombinase, was synthesized by gene synthesis and subcloned into the pUC57 vector (Genscript). LR recombination into the Gateway Destination vector containing Tol2 transposase sites and lens specific alpha-crystallin-dsRed sequence^44^ was performed using LR recombinase (ThermoFisher). Zebrafish embryos were coinjected with 25-40 pg of each construct and 75-150 pg of *Tol2* mRNA. Founders were screened for the presence of lens fluorescence in their progeny. Recombination activity was tested by crossing with *jam2b^Gt(2A-Gal4);^ Tg(actb2:BFP-loxP-dsRed-loxP)* driver / reporter line. One lines of each type, *UAS:Cre* and *UAS:CreERT2,* which showed good recombination activity, was selected and used for all subsequent experiments.

### Confocal microscopy imaging and image processing

For confocal imaging, embryos were anesthetized in 0.016% Tricaine solution, mounted in 0.6% low melting point agarose and imaged using a Nikon Eclipse or Nikon AX confocal microscope. A series of z-slices were acquired using NIS Elements software. The Nikon AI Denoise function was used to reduce the image noise. Maximum intensity projections were generated using Fiji or Nikon Elements software. Images were adjusted using the Levels function in Adobe Photoshop to enhance the brightness and contrast. In all cases, the same adjustments were applied to control and experimental embryo samples.

### Hybridization chain reaction (HCR)

Whole-mount fluorescent *in situ* hybridization was performed using the hybridization chain reaction version 3.0 as previously described.^45^ Fluorescently labeled RNA probes for *etv2*, *prrx1a, hand2, jam2a,* and *jam2b* were purchased from Molecular Instruments, Inc.

### Whole-mount *in situ* hybridization (ISH)

was performed as previously described.^46^ DIG-labeled antisense RNA probes for *jam2b,*^15^ *npas4l,*^15,47^ *etv2/etsrp,*^48^ *scl/tal1*^49^ were synthesized as previously described. To synthesize *Cre* probe, Cre DNA coding sequence was amplified by PCR from the plasmid template containing optimized Cre, which was used to generate UAS:Cre line. The following primers were used for PCR amplifcation: CATGCCCAAGAAAAAGCGTAAAGTG (forward) and CCTTTAATACGACTCACTATAGGGTTAGTCTCCATCCTCCAGCAGAC (reverse with T7 sequence attached). PCR product was purified using a spin column, and the DIG-labeled antisense probe was synthesized by in vitro transcription using T7 RNA polymerase (Promega). Following ISH, embryos were whole mounted in 0.6% low melting point agarose and imaged under light microscopy at 10× or 20× magnification with a Nikon Eclipse compound microscope. A series of z-slices were acquired using NIS Elements software to produce extended focus projection images.

### *jam2b* mRNA overexpression

The coding region of Jam2b was amplified by PCR from *jam2b* cDNA construct and subcloned into the pCSDest vector ^44^ using the two-way Gateway recombination approach (ThermoFisher). To synthesize mRNA for injections, the vector was linearized with NotI and transcribed with the SP6 mMessage mMachine Kit (ThermoFisher). 50-100 pg of mRNA was injected into embryos at the one-cell stage.

### 4-hydroxytamoxifen (OHT) treatments

To induce tissue-specific recombination at a specific time point, the *jam2b^Gt(2A-Gal4)^;UAS:CreERT2* line was crossed with the *Tg(actb2-loxP-BFP-loxP-dsRed; fli1:GFP)* reporter line. Embryos were treated with 10 µM 4-OHT (Sigma-Aldrich) solution starting at 7 or 22 hpf. Alternatively, to analyze the timing requirement, embryos were treated at 7 hpf stage and subsequently washed out at 22 hpf by rinsing several times in embryo water. Confocal z-stacks were acquired at 3 dpf and analyzed with NIS Elements Software. Recombination was quantified by counting dsRed-positive cells within the vasculature.

### Endothelial cell (EC) count

To assess EC numbers in *jam2a;jam2b* double homozygous mutants and control heterozygous embryos, confocal images were analyzed using NIS Elements Imaging Software. An region spanning 5 intersegmental vessels in the trunk region was selected, and ECs located in the floor of the posterior cardinal vein (PCV) were manually counted.

### Parachordal lymphangioblast count

Following the confocal imaging, 10 segments between adjacent intersegmental vessels from each control and mutant embryo were analyzed in NIS Elements. The fraction of horizontal myoseptum covered by PLs in each segment was estimated as 1/3, 2/3, or 3/3 (if PLs cover the entire segment). The average fraction of PL coverage over 10 segments was calculated and multiplied by 100% to obtain the percentage values.

### Subintestinal vessel and thoracic duct (TD) measurements

To evaluate SIA, SIV and TD in *jam2a; jam2b* double homozygous mutants and control heterozygous embryos, confocal images were analyzed using NIS Elements 5.41.02 Imaging Software. SIA, SIV and TD were categorized as normal or abnormal according to morphological characteristics. Vessels were considered normal if they were uniform and lacked gaps throughout the whole embryo; everything else was considered abnormal.

To analyze SIA formation in MZ *jam2a; jam2b* mutant and control embryos in *fli1:Kaede* background, the length of red and green only SIA segments was measured in Fiji (Image J), and the length percentage of red labeled SIA was calculated.

### Chemical cell ablation

To assess vascular tissue ablation and regeneration, *jam2a;jam2b* homozygous mutant zebrafish in a *kdrl:GFP* background were crossed with *jam2a;jam2b* homozygous mutants carrying the *etv2^Gt(2A-Gal4)^; UAS:NTR-mCherry* transgene. As a control group, heterozygous embryos were obtained by crossing *jam2a; jam2b* homozygous mutants carrying the *etv2^Gt(2A-Gal4)^; UAS:NTR-mCherry* transgene with wild-type *kdrl:GFP*. Embryos were collected and treated with 10 mM metronidazole (MTZ) dissolved in 0.1% DMSO at 6 or 10 hpf. At 2 dpf, embryos were dechorionated, thoroughly washed, and transferred to fresh embryo water to allow for recovery. Confocal imaging was performed at 3 dpf to assess vascular regeneration. NIS-Elements Imaging Software (version 5.41.02) was used to quantify tissue ablation and recovery. The total length of regenerated vessels was calculated by measuring and adding the lengths of all recovered vessels based on mCherry and GFP expression.

### Statistical analysis

Graphs were generated, and statistical analyses (t-tests and Fisher’s exact tests) were performed using GraphPad Prism software.

### Single-cell RNA-seq analysis

Transgenic *etv2^Gt(2A-Venus)^* fish were crossed to *jam2b^Gt(2A-Gal4)^;UAS:NTR-mCherry* fish. Double-positive embryos expressing both *Venus* and *mCherry* fluorescent proteins were selected at 36 hpf. Embryos were dechorionated and immediately transferred into a 1.5 ml Eppendorf tube to generate a single-cell suspension. Whole embryos were then dissociated into a single-cell suspension using a cold protease tissue dissociation protocol.^50^ Cells expressing *Venus*-only, *mCherry*-only, and both (*Venus* and *mCherry*) were sorted using a BD/FACSAria II flow cytometer cell sorter. After sorting, cells expressing both *Venus* and *mCherry* were divided equally and added to the *Venus*-only and *mCherry*-only populations manually. Suspensions of approximately 10,000 Venus-positive or mCherry-positive cells were loaded separately onto the 10X Genomics Single Cell 3’ (V3 chemistry) chip. 12 cDNA amplification cycles were used to generate cDNA. Samples were sequenced on an Illumina NovaSeq 6000 instrument (Illumina, San Diego, CA) running an S2 flow cell with parameters as follows: Read 1, 28 cycles; Index Read 1: 8 cycles; Read 2: 91 cycles at the Cincinnati Children’s Hospital Medical Center DNA Sequencing core. The fastq files obtained from the sequencing core were then mapped to the Danio rerio genome (version zv11) to generate single-cell feature counts using Cell Ranger version 3.0.2. Counts were generated using fastq files from each of the populations individually. Using the R packages: Seurat,^51^ ggplot2,^52^ dplyr,^53^ reshape2,^54^ and tidyverse,^55^ we filtered the data to retain high quality cells with 200-3500 features and less than 5% mitochondrial gene content. The datasets were then integrated and the plots were generated using custom written R code.^56-58^ To identify a stable cluster resolution, we tested resolutions from 0 to 3 in 0.1 increments and selected 1.6, which revealed the SVF cells in cluster 15. We then selected clusters for trajectory analysis using the Monocle 3 package. The UMAP, feature plots, and violin plots were generated using Seurat, and the lineage plot was generated using Monocle 3.^26^ Cluster annotation was done manually based on the expression of top marker genes. In some cases, assistance from ChatGPT 5 was used to for cell type prediction based on the 40-60 top marker genes, which was subsequently verified manually based on published gene expression patterns. The single-cell RNA-seq dataset has been deposited to the NCBI GEO database under accession number: GSE179926.

## Acknowledgements

This research was supported by NIH R01 HL153005 to S.S., a supplemental postdoctoral diversity award R01 HL163161-S1 to R.D., AHA 19POST34400016 award to S.M., AHA 19POST34450021 award to S.G., and AHA 26POST1551571 award to S.P.R.B. We thank Olivia Nester, Kanwal Jussa and Melanie Taylor for their assistance with zebrafish experiments.

---

**Supplementary Figure S1.**
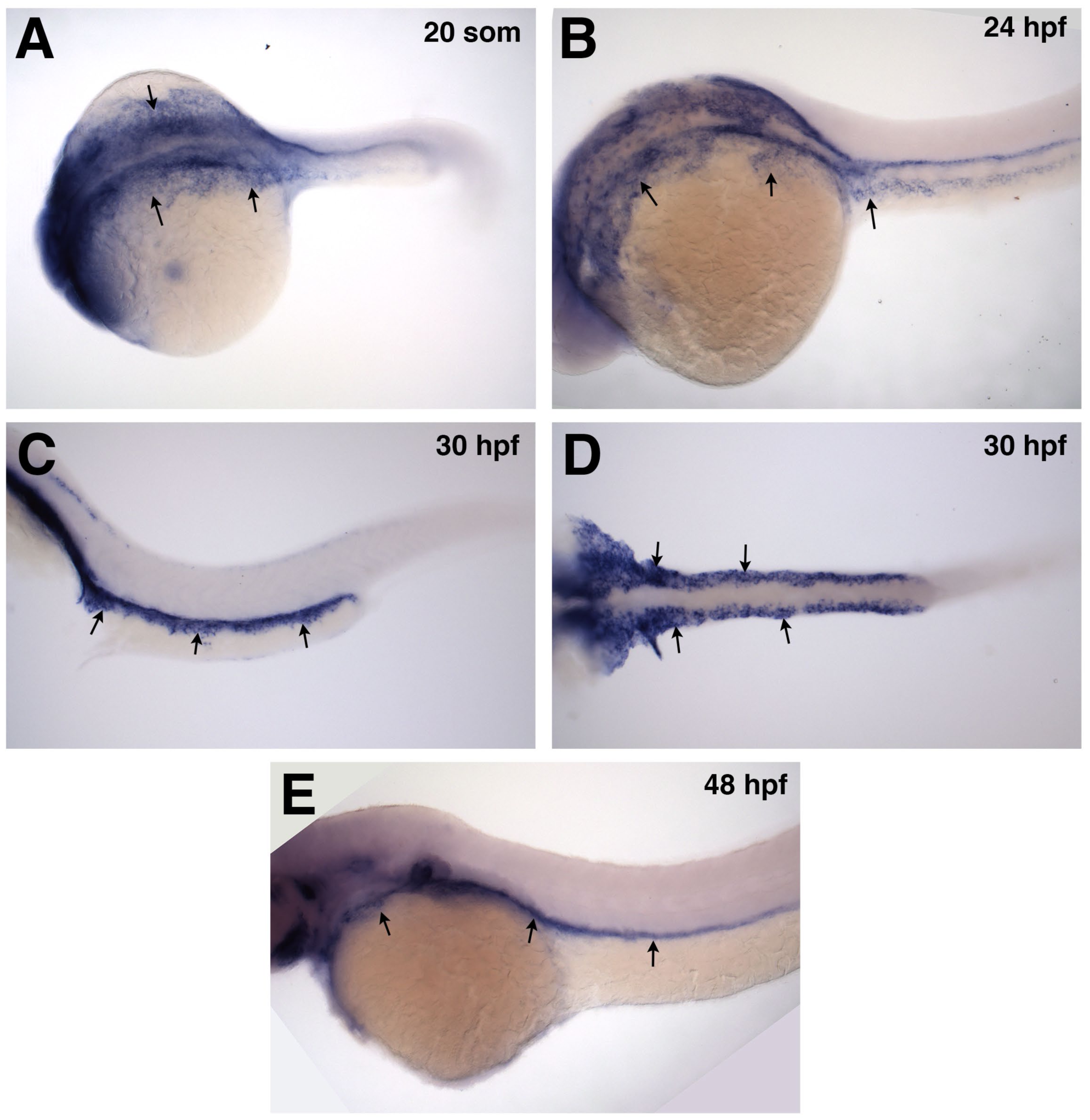
In situ hybridization analysis of *jam2b* expression between the 20-somite and 48 hpf stages. Note strong *jam2b* expression in the lateral plate mesoderm along the yolk and yolk extension (arrows). (A,B,C,E) lateral views, (D) ventral view.

**Supplementary Figure S2.**
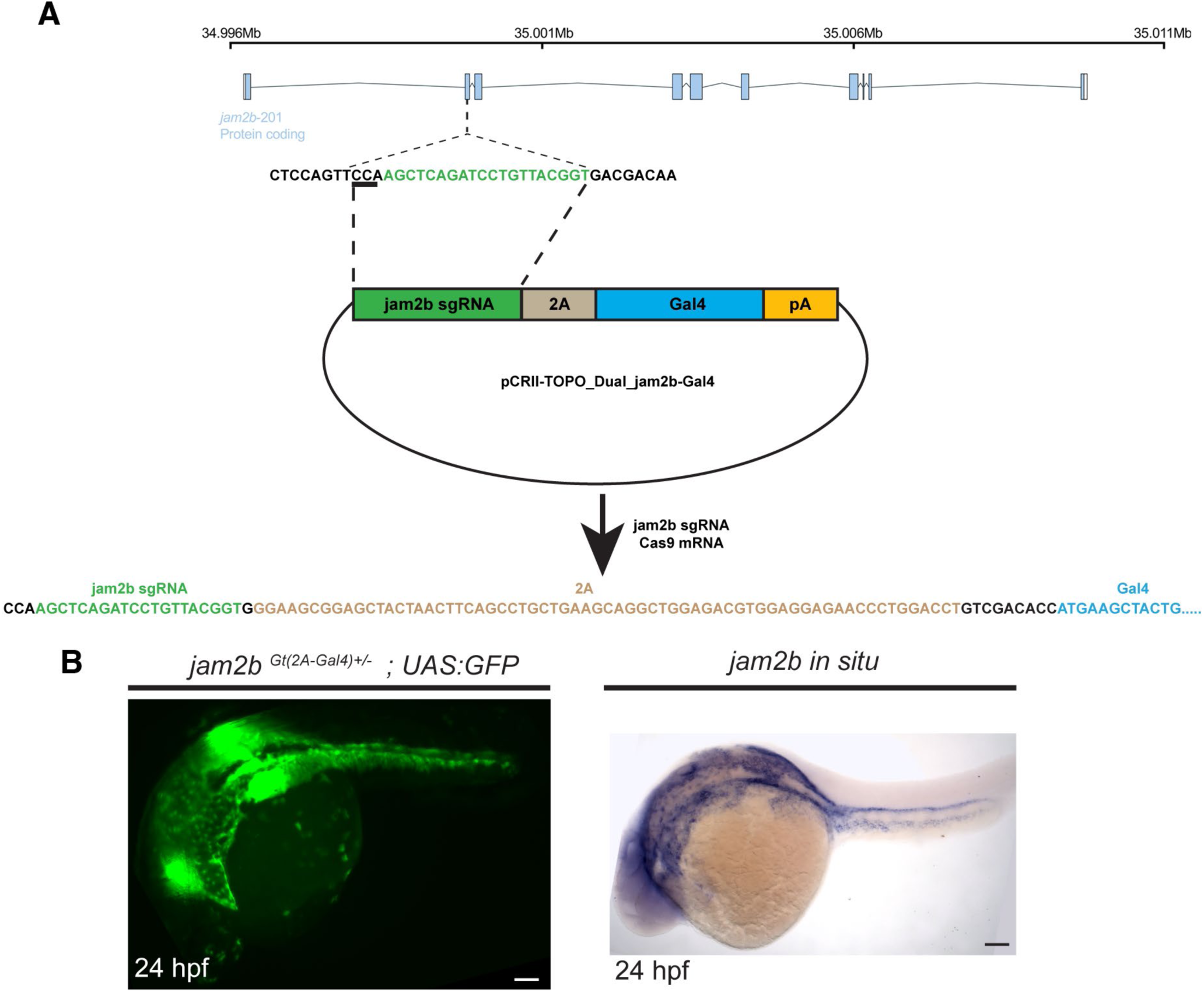
Generation of *jam2b^Gt(2A-Gal4)^; UAS:GFP* line using CRISPR-Cas9 mediated non-homologous recombination. (A) A diagram of *sgRNA-2A-Gal4* construct and its insertion into the *jam2b* genomic locus. The targeting construct contained the *jam2b* sgRNA site, followed by an in-frame fusion to the viral P2A peptide, the Gal4 DNA-binding domain, and the SV40 polyA sequence. The *jam2b* sgRNA targets the second exon of the *jam2b* genomic sequence. (B) Comparison of GFP fluorescence in a *jam2b^Gt(2A-Gal4)^; UAS:GFP* embryo with *jam2b* mRNA expression analyzed by in situ hybridization at 24 hpf. Note bilateral *jam2b* expression and GFP fluorescence observed in the lateral mesoderm along the yolk and yolk extension. Scale bars: 100 µm.

**Supplementary Figure S3.**
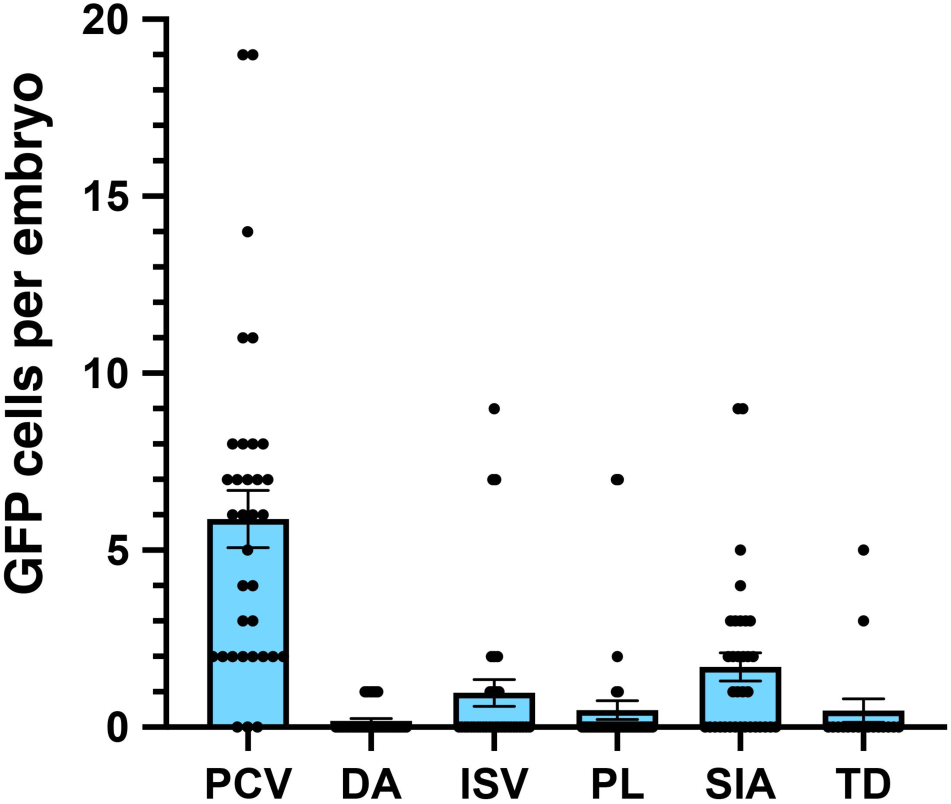
**Quantification of *jam2b^Gt(2A-Gal4)^; UAS:GFP* cell contribution to different vascular beds**. All analyses were performed at 3 dpf, except for TD, which is at 4 dpf. n=34 embryos, except for TD, for which n=17. PCV, posterior cardinal vein, DA, dorsal aorta, ISV, intersegmental vessel, PL, parachordal lymphangioblast, SIA, supraintestinal artery, TD, thoracic duct.

**Supplementary Figure S4.**
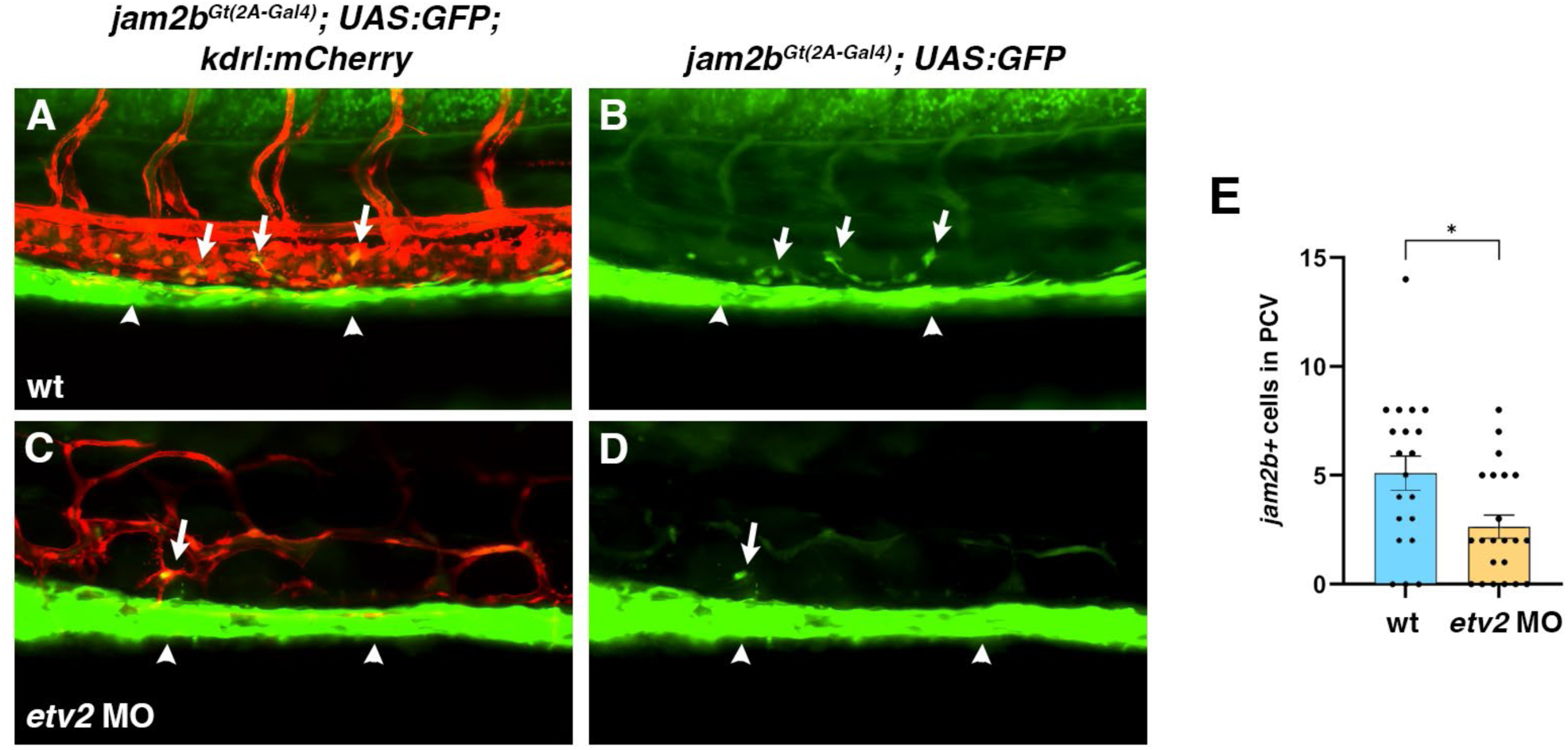
Contribution of *jam2b+* cells to the PCV depends on Etv2 function. (A,B) Confocal microscopy analysis of GFP+ cells in the PCV (arrows) of *jam2b^Gt(2A-Gal4)^; UAS:GFP; kdrl:mCherry* embryos at 72 hpf. (C,D) *Etv2* MO-injected embryos show reduced contribution of GFP+ cells to the PCV (arrows), while GFP expression in the LPM (arrowheads) is not affected. In addition, ISV sprouting and vascular patterning are greatly perturbed by *etv2* knockdown. (E) Quantification of GFP+ cells in the PCV in uninjected (n=20) and *etv2* MO-injected (n=23) embryos. Outliers were removed using Grubbs’ test (alpha=0.05) and Welch’s t-test was used for statistical analysis, *p<0.05.

**Supplementary Figure S5.**
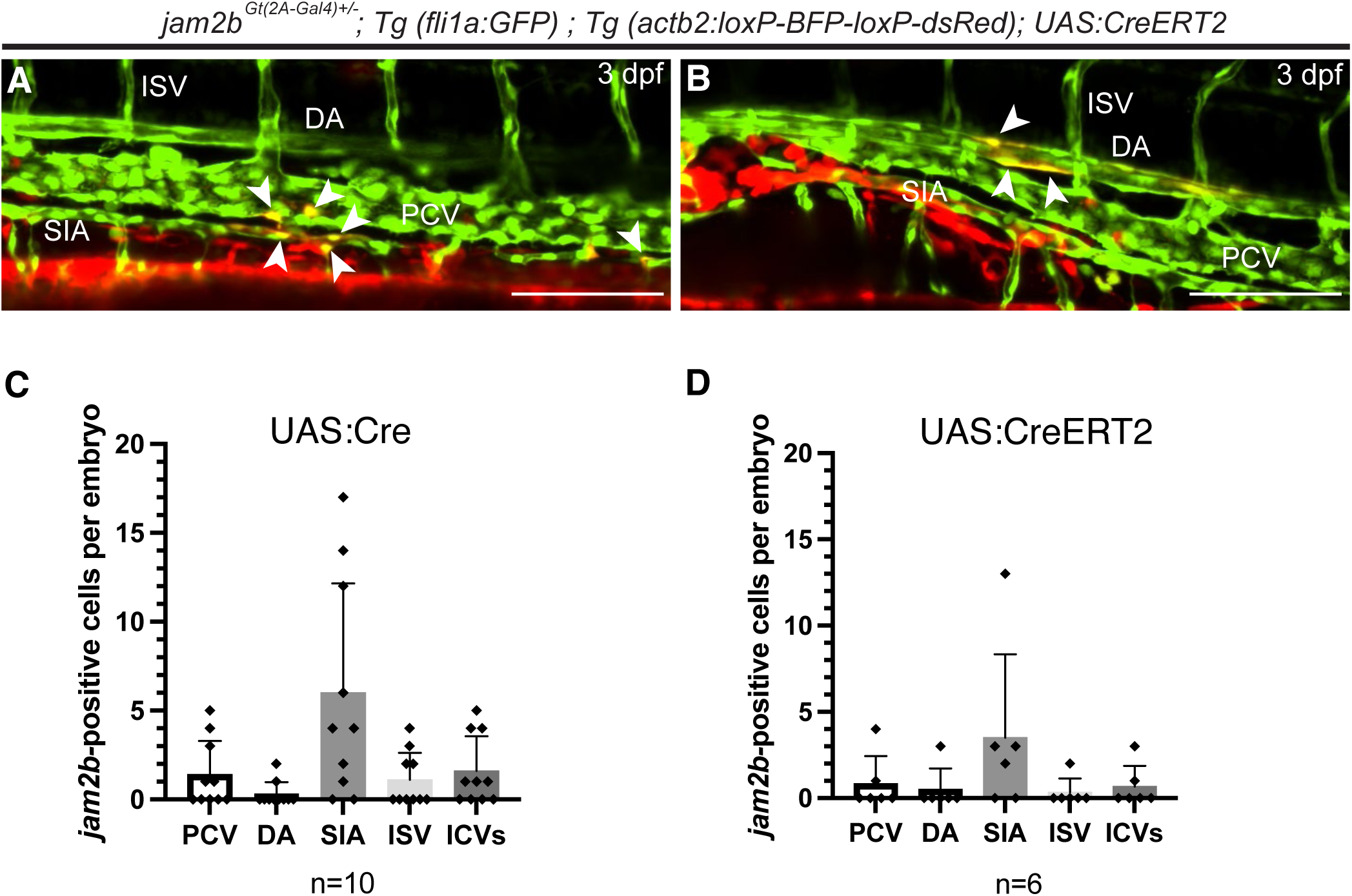
Fate mapping of *jam2b+* positive cells at 3 dpf based on transient *UAS:Cre* and *UAS:CreERT2* expression. Plasmids containing *UAS:Cre* or *UAS:CreERT2* were co-injected with *Tol2* mRNA into *jam2b^Gt(2A-Gal4)+/-^; fli1:GFP; actb2:loxP-BFP-loxP-dsRed* embryos. Embryos were imaged by confocal microscopy at 3 dpf. 4-OHT was added at 7 hpf to embryos injected with *UAS:CreERT2*. Note the presence of switched dsRed-positive cells in the SIA and PCV (arrowheads, A), and the DA (B). (C,D) Quantification of dsRed cell contribution to different vascular beds. PCV, posterior cardinal vein; DA, dorsal aorta; SIA, supraintestinal artery; ISV, intersegmental vessel, ICVs, interconnecting vessels along the yolk and yolk extension. Lateral view of the trunk region; anterior is to the left. Scale bars: 100 μm.

**Supplementary Figure S6.**
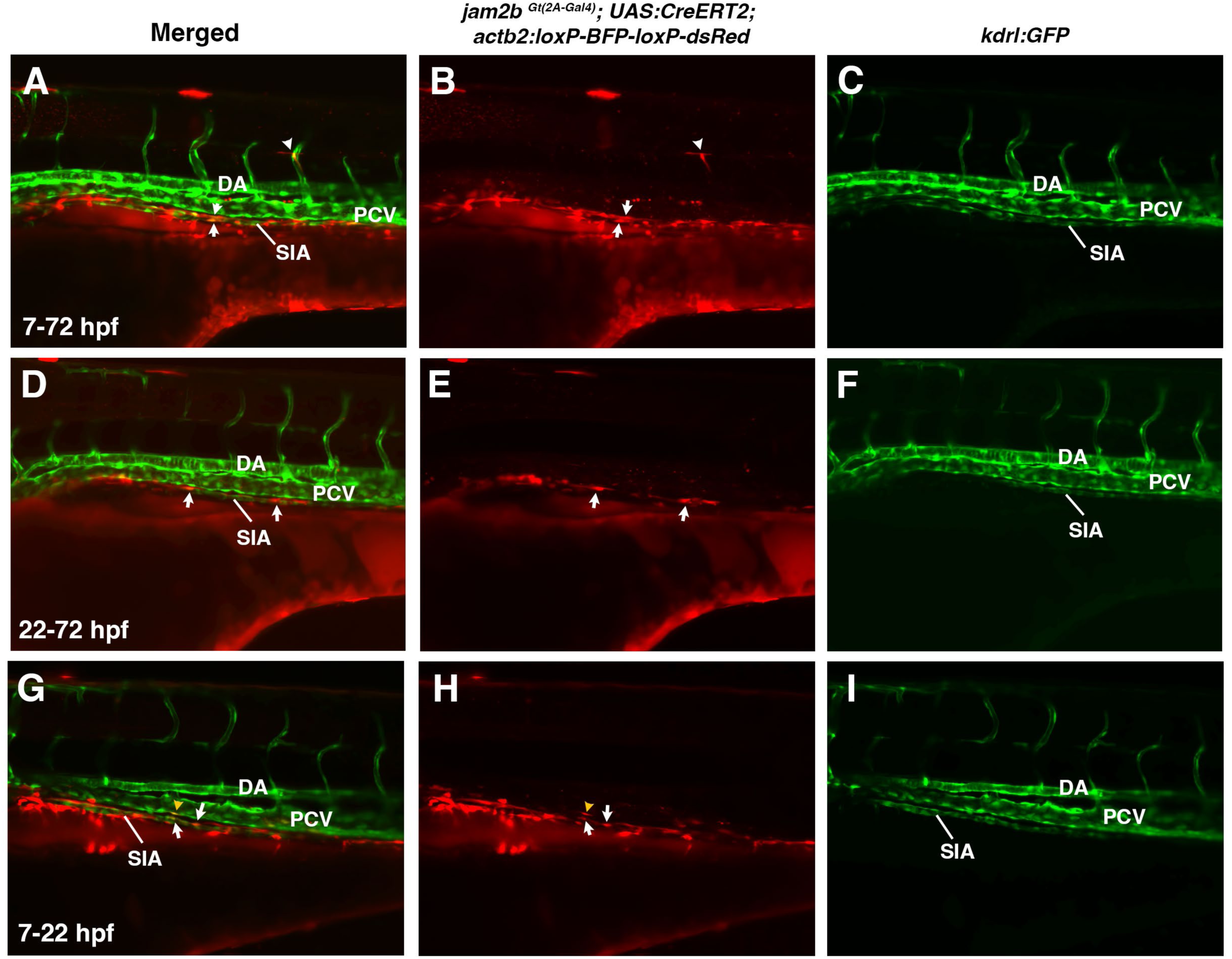
Fate mapping of endothelial cell labeling in different vascular beds in *jam2b^Gt(2A-Gal4)^; UAS:CreERT2; actb2-loxP-BFP-loxP-dsRed* embryos at 3 dpf. (A-C) 4-OHT was added starting at 7 hpf; (D-F) 4-OHT was added at 22 hpf. (G-I) 4-OHT was added at 7 hpf and washed out at 22 hpf stages. Note the frequent *jam2b+* cell contribution to the SIA (white arrows), and less frequent contribution to parachordal lymphangioblasts (white arrowheads, A, B) and the PCV (yellow arrowheads, G, H). In addition, many dsRed+ cells are located in the mesothelial layer over the yolk extension. Cell contribution to the SIV could not be analyzed reliably due to very strong red fluorescence in the mesothelial layer surrounding the yolk.

**Supplementary Figure S7.**
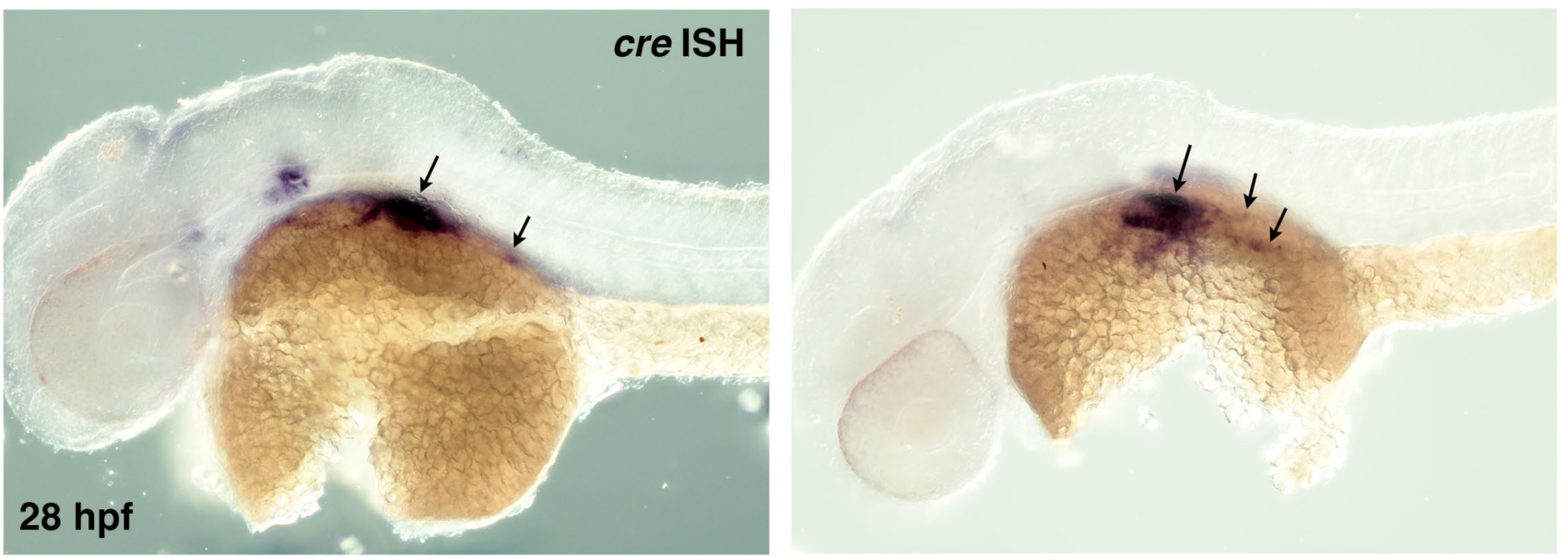
In situ hybridization analysis for Cre mRNA expression in *jam2b^Gt(2A-Gal4)^; UAS:Cre* embryos at 28 hpf. Note partial and fragmented expression of Cre in the lateral plate mesoderm adjacent to the yolk, indicative of a mosaic and partially silenced expression pattern.

**Supplementary Figure S8.**
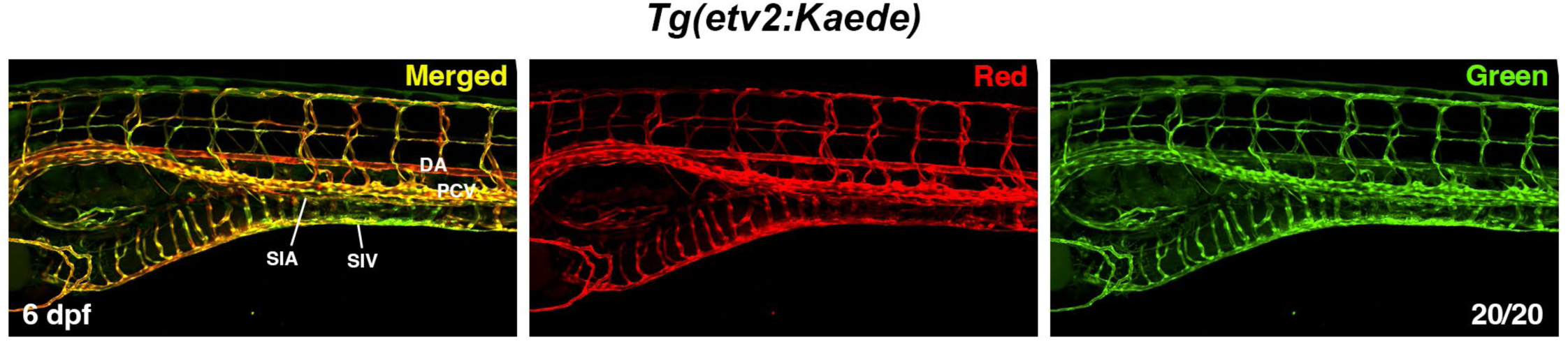
No new progenitors contribute to intestinal vasculature between 3 and 6 dpf. *Tg(etv2:Kaede)* larvae were photoconverted at 3 dpf and then imaged at 6 dpf by confocal microscopy. The SIA and SIV are positive for both green and red.

**Supplementary Figure S9.**
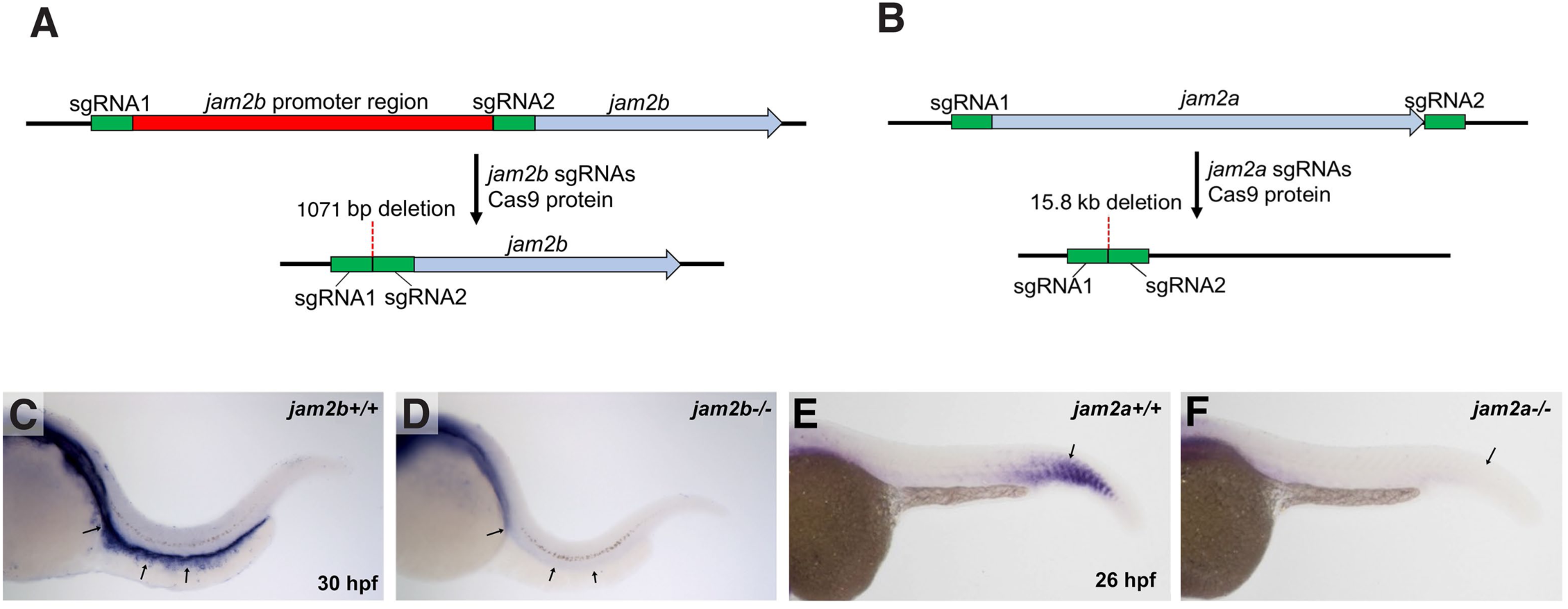
Generation of *jam2b* and *jam2a* mutants using CRISPR-Cas9 genome editing. (A) A schematic diagram illustrating generation of *jam2b* mutants using two sgRNAs designed to delete the promoter region. (B) A schematic diagram illustrating generation of *jam2a* mutants using two sgRNAs designed to delete the entire *jam2a* coding sequence. (C,D) In situ hybridization (ISH) analysis for *jam2b* expression in wild-type *jam2b+/+* and *jam2b-/-* mutant embryos at 30 hpf. Note that *jam2b* expression along the lateral plate mesoderm is absent in *jam2b* mutants (arrows). Some residual staining corresponds to background signal and pigment cells along the horizontal myoseptum. (E,F) ISH analysis for *jam2a* expression in wild-type *jam2a+/+* and *jam2a-/-* mutants at 26 hpf. The strongest *jam2a* expression is observed in the posterior somites, which is absent in *jam2a* mutants (arrows).

**Supplementary Figure S10.**
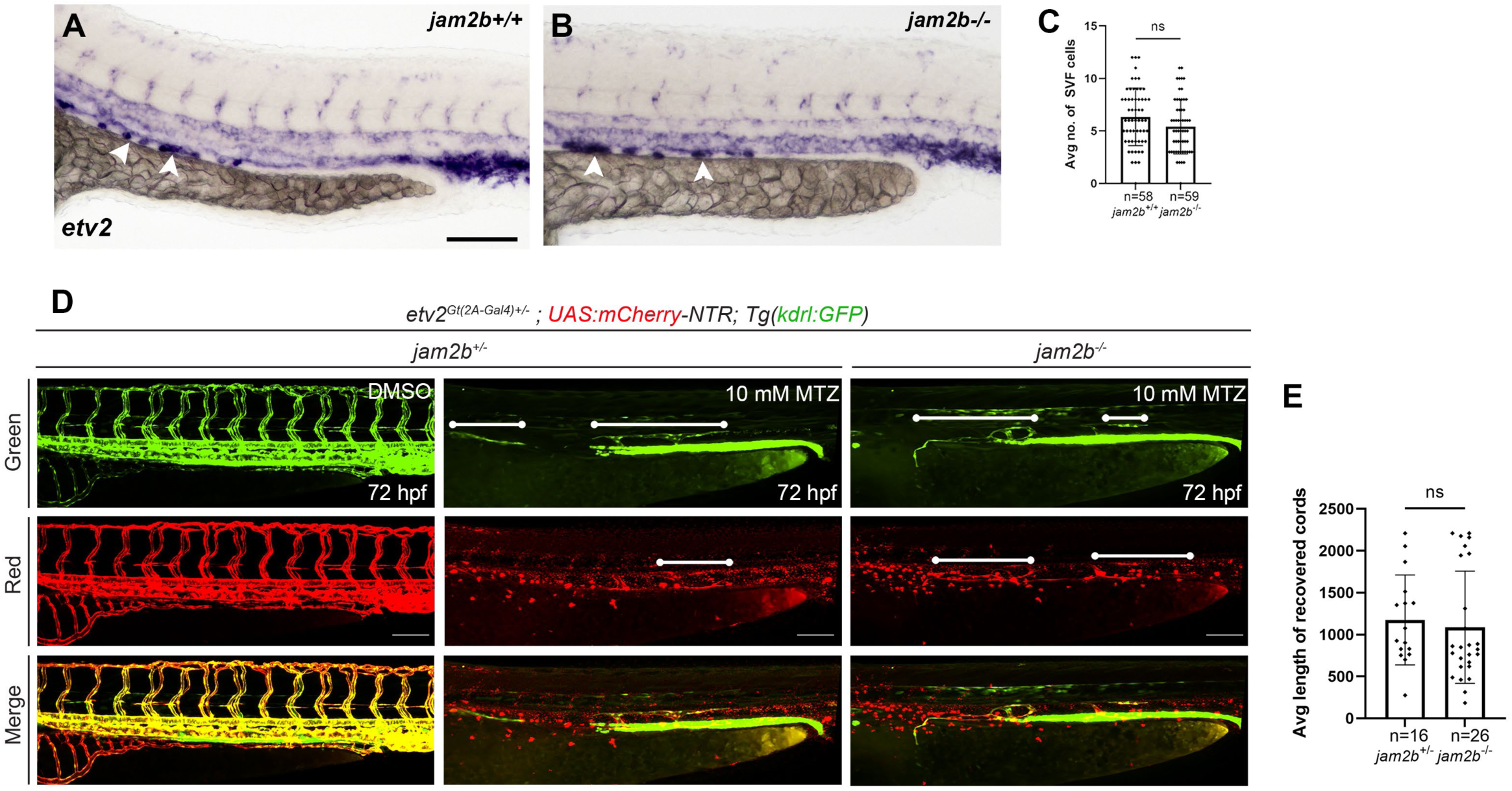
***jam2b* mutant embryos do not show any apparent defects in SVF cell formation or vascular recovery after endothelial cell ablation.** (A-C) Quantification of SVF cells (white arrowheads) by in situ hybridization analysis for *etv2* expression at 30 hpf. Experimental and control embryos were obtained from the incrosses of *jam2b-/-* and wild-type (*jam2b+/+*) sibling parents in a *kdrl:GFP* background, respectively. (D,E) Quantification of vascular recovery after endothelial cell ablation. Embryos were obtained by crossing *jam2b+/-* and *jam2b-/-* adults in *etv2^Gt(2A-Gal4)^; UAS:mCherry-NTR; Tg(kdrl:GFP)* background were crossed with each other. Embryos were treated with 10 mM MTZ from 10 hpf to 45 hpf, after which MTZ was washed out and the embryos were allowed to undergo vascular recovery until 72 hpf, when they were imaged and genotyped. Control embryos were treated with 0.1% DMSO. The length of new vascular cords (white lines) at the position of the PCV and SIV was measured and compared in *jam2b+/-* and *jam2b-/-* mutant embryos. Note that the bright *kdrl:GFP* expression along the yolk extension corresponds to the pronephros and was not affected in these experiments. Lateral view is shown; anterior is to the left. Mean ± s.d. is shown; ns, not significant. Scale bars: 100 μm.

**Supplementary Figure S11.**
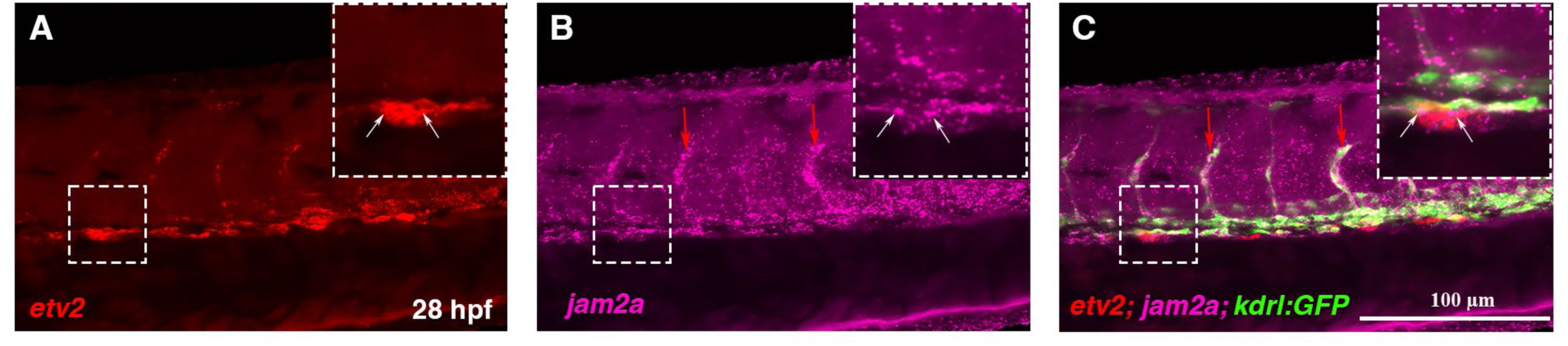
Expression of *jam2a* at 28 hpf. (A) *etv2* (red) and (B) *jam2a* mRNA (magenta) expression detected by HCR. White arrows in the enlarged panel indicate *jam2a* coexpression with *etv2* in SVF cells. (C) Merged image showing *jam2a* and *etv2* expression in the *kdrl:GFP* line. Note *jam2a* expression in perivascular cells adjacent to the axial and intersegmental vessels (red arrows).

**Supplementary Figure S12.**
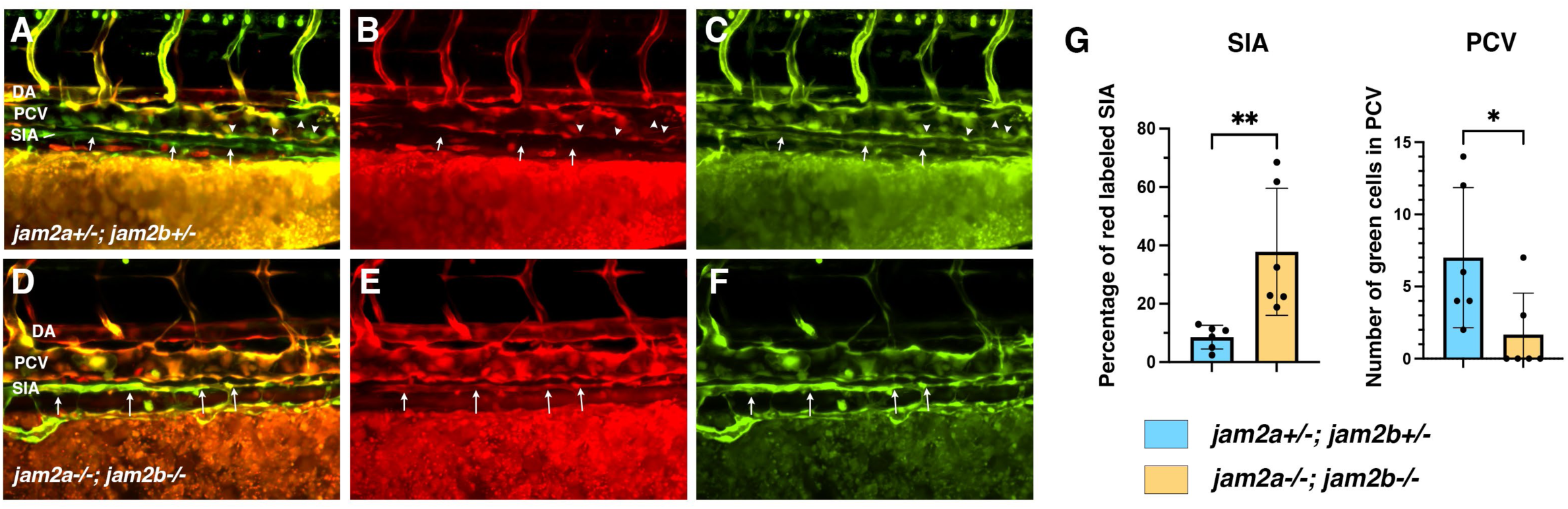
Analysis of new vasculogenesis in *jam2a; jam2b; fli1:Kaede* embryos at 3 dpf. MZ *jam2a-/-; jam2b-/-* embryos and control embryos in *fli1:Kaede* background were photoconverted at 24 hpf and then analyzed by confocal imaging at 3 dpf. Control embryos were obtained by the cross of *jam2a+/+; jam2b+/-* and *jam2b-/-; jam2b+/-* parents and thus represent a mixture of *jam2a+/-; jam2b±/±* genotypes. (A-F) Mid-trunk region of merged (A,D), red (B,E) and green (C,F) channels is shown. Note that the SIA in control embryos is largely green-only (arrows, A-C), whereas large sections of SIA in MZ *jam2a; jam2b* mutants show red fluorescence (arrows, D-F), indicating that it has formed from endothelial cells that were present before 24 hpf. Green only cells, which formed by new vasculogenesis, are apparent in the PCV of control embryos (arrowheads, A-C), while they are largely absent in MZ *jam2a; jam2b* mutants. (G) Quantification of the percentage of the SIA, labeled by red fluorescence (left graph) and the number of green-only cells in the PCV of control and MZ *jam2a-/-; jam2b-/-* embryos. **p<0.01, *p<0.05, two-tailed t-test.

**Supplementary Figure S13.**
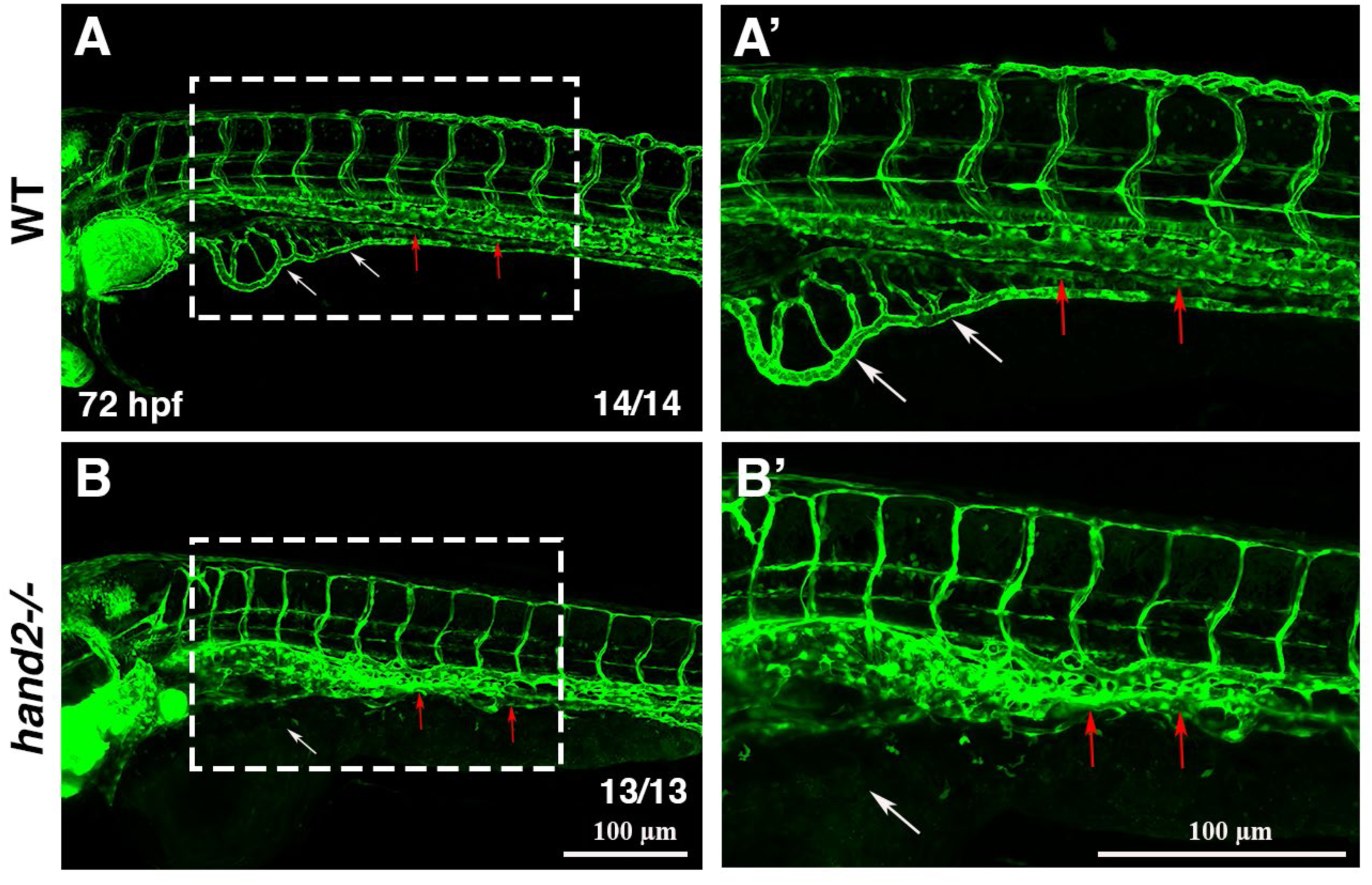
Absence of intestinal vasculature in *hand2* mutant embryos at 72 hpf. (A) Representative wild-type (WT) embryo showing the intestinal vasculature visualized using the *fli1:GFP* line. White arrows indicate the subintestinal vein (SIV), and red arrows indicate the supraintestinal artery (SIA). (A’) Enlarged 20× view of the boxed region in (A). (B) Representative *hand2* mutant embryo lacking detectable SIV and SIA, indicated by white and red arrows, respectively. In addition, the DA and PCV fail to separate. (B’) Enlarged 20× view of the boxed region in (B).

**Supplementary Figure S14.**
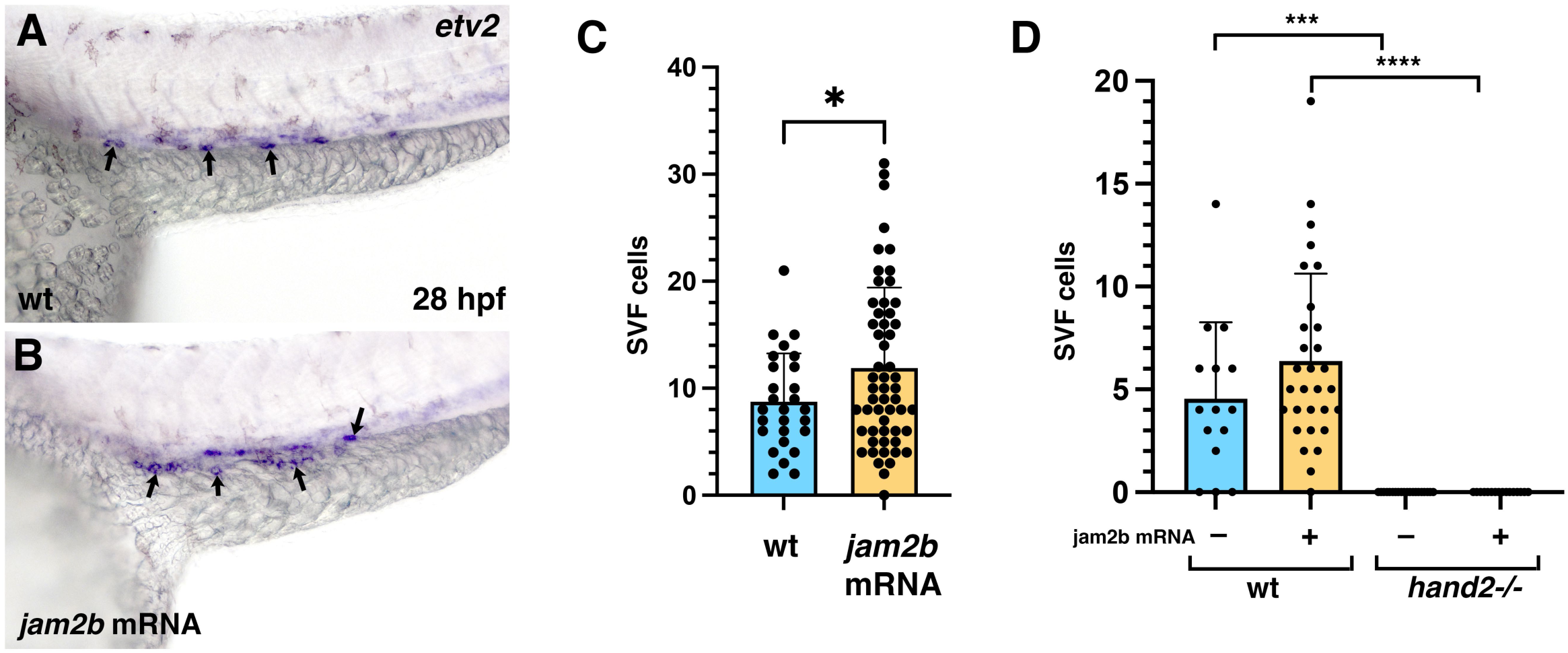
***jam2b* overexpression expands SVF cell formation in the presence of *hand2* function.** (A,B) *etv2* expression analysis in SVF cells (arrows) by in situ hybridization in uninjected wild-type embryos and embryos injected with 100 pg of *jam2b* mRNA at 28 hpf. Note the expansion of SVF cells along the yolk extension in *jam2b* mRNA injected embryos. (C) Quantification of *etv2+* SVF cells in uninjected control (wt) and *jam2b* mRNA injected embryos at 28 hpf. p<0.05, two-tailed t-test. (D) *jam2b* mRNA fails to restore *etv2+* SVF cells in *hand2* mutants at 26 hpf. Quantification of SVF cells in wt uninjected (n=15), wt injected with 100 pg *jam2b* mRNA (n=30), *hand2* mutants uninjected (n=20), and *hand2* mutants injected with 100 pg *jam2b* mRNA (n=16). Note that the number of SVF cells was 0 in all *hand2* mutant samples. Embryos from an incross of *hand2+/-* adults in a *myl7:GFP* background were injected with *jam2b* mRNA, and homozygous mutants and wild-type siblings were sorted at 24-26 hpf based on *myl7:GFP* fluorescence pattern; the wild-type group included both *hand2+/+* and *hand2+/-* embryos. ***p=0.0001; ****p<0.0001, one-way ANOVA followed by Šidák’s multiple comparisons test.

## MOVIE LEGENDS

**Movie 1. Migration of *jam2b*-expressing cells from the lateral plate mesoderm into the PCV.** A time-lapse video of a *jam2b^Gt(2A-Gal4)^;UAS:GFP;Tg(kdrl:mCherry)* embryo shows GFP positive cells migrating dorsally, integrating into the PCV, proliferating and initiating *kdrl:mCherry* expression (white arrowheads) (see Fig. 2A-F). Imaging was started at 24 hpf. PCV, posterior cardinal vein; DA, dorsal aorta. Scale bar: 20 µm.

**Movie 2. Migration of *jam2b*-expressing cells from the lateral plate mesoderm into the ISV.** A time-lapse video of a *jam2b^Gt(2A-Gal4)^;UAS:GFP;Tg(kdrl:mCherry)* embryo shows GFP positive cells migrating into venous ISV (white arrow). Another GFP+ cell can be seen, which remains in the PCV (white arrowhead). Imaging was started at 24 hpf.

**Movie 3. Contribution of *jam2b*-expressing cells to the parachordal lymphangioblasts (PLs).** A time-lapse video of a *jam2b^Gt(2A-Gal4)^;UAS:GFP;Tg(kdrl:mCherry)* embryo shows a GFP positive cell migrating out of the LPM and contributing to the PLs at the horizontal myoseptum (white arrowhead). An adjacent skeletal muscle cell with strong GFP expression is apparent next to the GFP-positive PL. Imaging was started at 24 hpf.

**Movie 4. *jam2b*-expressing cells are derived from the lateral plate mesoderm:** A time-lapse video of a *jam2b^Gt(2A-Gal4)^;UAS:GFP;Tg(kdrl:mCherry)* embryo showing from the 18-somite stage to 24 hpf. GFP positive cells can be seen in the ventrolateral LPM region at the 18-somite stage and remain in the same region until 24 hpf as the embryo undergoes convergent extension. PCV, posterior cardinal vein; DA, dorsal aorta. Scale bar: 50 µm.

## REFERENCES

1. Gouysse, G., Couvelard, A., Frachon, S., Bouvier, R., Nejjari, M., Dauge, M.C., Feldmann, G., Henin, D., and Scoazec, J.Y. (2002). Relationship between vascular development and vascular differentiation during liver organogenesis in humans. J Hepatol 37, 730–740. 10.1016/s0168-8278(02)00282-9.

2. Kingsbury, J.W., Alexanderson, M., and Kornstein, E.S. (1956). The development of the liver in the chick. Anat Rec 124, 165–187. 10.1002/ar.1091240204.

3. Matsumoto, K., Yoshitomi, H., Rossant, J., and Zaret, K.S. (2001). Liver organogenesis promoted by endothelial cells prior to vascular function. Science 294, 559–563. 10.1126/science.1063889.

4. Sherer, G.K. (1991). The Development of the Vascular System (Karger).

5. Goldman, O., Han, S., Hamou, W., Jodon de Villeroche, V., Uzan, G., Lickert, H., and Gouon-Evans, V. (2014). Endoderm generates endothelial cells during liver development. Stem Cell Reports 3, 556–565. 10.1016/j.stemcr.2014.08.009.

6. Perez-Pomares, J.M., Carmona, R., Gonzalez-Iriarte, M., Macias, D., Guadix, J.A., and Munoz-Chapuli, R. (2004). Contribution of mesothelium-derived cells to liver sinusoids in avian embryos. Dev Dyn 229, 465–474. 10.1002/dvdy.10455.

7. Zhang, H., Pu, W., Tian, X., Huang, X., He, L., Liu, Q., Li, Y., Zhang, L., He, L., Liu, K., et al. (2016). Genetic lineage tracing identifies endocardial origin of liver vasculature. Nat Genet 48, 537–543. 10.1038/ng.3536.

8. Koenig, A.L., Baltrunaite, K., Bower, N.I., Rossi, A., Stainier, D.Y., Hogan, B.M., and Sumanas, S. (2016). Vegfa signaling promotes zebrafish intestinal vasculature development through endothelial cell migration from the posterior cardinal vein. Dev Biol 411, 115–127. 10.1016/j.ydbio.2016.01.002.

9. Goi, M., and Childs, S.J. (2016). Patterning mechanisms of the sub-intestinal venous plexus in zebrafish. Dev Biol 409, 114–128. 10.1016/j.ydbio.2015.10.017.

10. Hen, G., Nicenboim, J., Mayseless, O., Asaf, L., Shin, M., Busolin, G., Hofi, R., Almog, G., Tiso, N., Lawson, N.D., and Yaniv, K. (2015). Venous-derived angioblasts generate organ-specific vessels during zebrafish embryonic development. Development 142, 4266–4278. 10.1242/dev.129247.

11. Eriksson, J., and Lofberg, J. (2000). Development of the hypochord and dorsal aorta in the zebrafish embryo (Danio rerio). J Morphol 244, 167–176. 10.1002/(SICI)1097-4687(200006)244:3<167::AID-JMOR2>3.0.CO;2-J.

12. Fouquet, B., Weinstein, B.M., Serluca, F.C., and Fishman, M.C. (1997). Vessel patterning in the embryo of the zebrafish: guidance by notochord. Dev Biol 183, 37–48. 10.1006/dbio.1996.8495.

13. Childs, S., Chen, J.N., Garrity, D.M., and Fishman, M.C. (2002). Patterning of angiogenesis in the zebrafish embryo. Development 129, 973–982. 10.1242/dev.129.4.973.

14. Jin, S.W., Beis, D., Mitchell, T., Chen, J.N., and Stainier, D.Y. (2005). Cellular and molecular analyses of vascular tube and lumen formation in zebrafish. Development 132, 5199–5209. 10.1242/dev.02087.

15. Metikala, S., Warkala, M., Casie Chetty, S., Chestnut, B., Rufin Florat, D., Plender, E., Nester, O., Koenig, A.L., Astrof, S., and Sumanas, S. (2022). Integration of vascular progenitors into functional blood vessels represents a distinct mechanism of vascular growth. Dev Cell 57, 767–782 e766. 10.1016/j.devcel.2022.02.015.

16. Metikala, S., Casie Chetty, S., and Sumanas, S. (2021). Single-cell transcriptome analysis of the zebrafish embryonic trunk. PLoS One 16, e0254024. 10.1371/journal.pone.0254024.

17. Ebnet, K. (2017). Junctional Adhesion Molecules (JAMs): Cell Adhesion Receptors With Pleiotropic Functions in Cell Physiology and Development. Physiol Rev 97, 1529–1554. 10.1152/physrev.00004.2017.

18. Kobayashi, I., Kobayashi-Sun, J., Hirakawa, Y., Ouchi, M., Yasuda, K., Kamei, H., Fukuhara, S., and Yamaguchi, M. (2020). Dual role of Jam3b in early hematopoietic and vascular development. Development 147. 10.1242/dev.181040.

19. Kobayashi, I., Kobayashi-Sun, J., Kim, A.D., Pouget, C., Fujita, N., Suda, T., and Traver, D. (2014). Jam1a-Jam2a interactions regulate haematopoietic stem cell fate through Notch signalling. Nature 512, 319–323. 10.1038/nature13623.

20. Powell, G.T., and Wright, G.J. (2012). Genomic organisation, embryonic expression and biochemical interactions of the zebrafish junctional adhesion molecule family of receptors. PLoS One 7, e40810. 10.1371/journal.pone.0040810.

21. Thisse, B., and Thisse, C. (2004). Fast Release Clones: A High Throughput Expression Analysis.

22. Auer, T.O., Duroure, K., De Cian, A., Concordet, J.P., and Del Bene, F. (2014). Highly efficient CRISPR/Cas9-mediated knock-in in zebrafish by homology-independent DNA repair. Genome Res 24, 142–153. 10.1101/gr.161638.113.

23. Chestnut, B., Casie Chetty, S., Koenig, A.L., and Sumanas, S. (2020). Single-cell transcriptomic analysis identifies the conversion of zebrafish Etv2-deficient vascular progenitors into skeletal muscle. Nat Commun 11, 2796. 10.1038/s41467-020-16515-y.

24. Akitake, C.M., Macurak, M., Halpern, M.E., and Goll, M.G. (2011). Transgenerational analysis of transcriptional silencing in zebrafish. Dev Biol 352, 191–201. 10.1016/j.ydbio.2011.01.002.

25. Kohli, V., Schumacher, J.A., Desai, S.P., Rehn, K., and Sumanas, S. (2013). Arterial and venous progenitors of the major axial vessels originate at distinct locations. Dev Cell 25, 196–206. 10.1016/j.devcel.2013.03.017.

26. Cao, J., Spielmann, M., Qiu, X., Huang, X., Ibrahim, D.M., Hill, A.J., Zhang, F., Mundlos, S., Christiansen, L., Steemers, F.J., et al. (2019). The single-cell transcriptional landscape of mammalian organogenesis. Nature 566, 496–502. 10.1038/s41586-019-0969-x.

27. Powell, G.T., and Wright, G.J. (2011). Jamb and jamc are essential for vertebrate myocyte fusion. PLoS Biol 9, e1001216. 10.1371/journal.pbio.1001216.

28. Yelon, D., Ticho, B., Halpern, M.E., Ruvinsky, I., Ho, R.K., Silver, L.M., and Stainier, D.Y. (2000). The bHLH transcription factor hand2 plays parallel roles in zebrafish heart and pectoral fin development. Development 127, 2573–2582. 10.1242/dev.127.12.2573.

29. Prummel, K.D., Crowell, H.L., Nieuwenhuize, S., Brombacher, E.C., Daetwyler, S., Soneson, C., Kresoja-Rakic, J., Kocere, A., Ronner, M., Ernst, A., et al. (2022). Hand2 delineates mesothelium progenitors and is reactivated in mesothelioma. Nat Commun 13, 1677. 10.1038/s41467-022-29311-7.

30. Ming, Z., Liu, F., Moran, H.R., Lalonde, R.L., Adams, M., Restrepo, N.K., Joshi, P., Ekker, S.C., Clark, K.J., Friedberg, I., et al. (2025). Lineage labeling with zebrafish hand2 Cre and CreERT2 recombinase CRISPR knock-ins. Dev Dyn. 10.1002/dvdy.70022.

31. Perens, E.A., Garavito-Aguilar, Z.V., Guio-Vega, G.P., Pena, K.T., Schindler, Y.L., and Yelon, D. (2016). Hand2 inhibits kidney specification while promoting vein formation within the posterior mesoderm. Elife 5. 10.7554/eLife.19941.

32. Meguenani, M., Miljkovic-Licina, M., Fagiani, E., Ropraz, P., Hammel, P., Aurrand-Lions, M., Adams, R.H., Christofori, G., Imhof, B.A., and Garrido-Urbani, S. (2015). Junctional adhesion molecule B interferes with angiogenic VEGF/VEGFR2 signaling. FASEB J 29, 3411–3425. 10.1096/fj.15-270223.

33. Casie Chetty, S., Rost, M.S., Enriquez, J.R., Schumacher, J.A., Baltrunaite, K., Rossi, A., Stainier, D.Y., and Sumanas, S. (2017). Vegf signaling promotes vascular endothelial differentiation by modulating etv2 expression. Dev Biol 424, 147–161. 10.1016/j.ydbio.2017.03.005.

34. Perens, E.A., and Yelon, D. (2025). Drivers of vessel progenitor fate define intermediate mesoderm dimensions by inhibiting kidney progenitor specification. Dev Biol 517, 126–139. 10.1016/j.ydbio.2024.09.008.

35. Proulx, K., Lu, A., and Sumanas, S. (2010). Cranial vasculature in zebrafish forms by angioblast cluster-derived angiogenesis. Dev Biol 348, 34–46. 10.1016/j.ydbio.2010.08.036.

36. Lawson, N.D., and Weinstein, B.M. (2002). In vivo imaging of embryonic vascular development using transgenic zebrafish. Dev Biol 248, 307–318. 10.1006/dbio.2002.0711.

37. Chestnut, B., and Sumanas, S. (2020). Zebrafish etv2 knock-in line labels vascular endothelial and blood progenitor cells. Dev Dyn 249, 245–261. 10.1002/dvdy.130.

38. Bertrand, J.Y., Chi, N.C., Santoso, B., Teng, S., Stainier, D.Y., and Traver, D. (2010). Haematopoietic stem cells derive directly from aortic endothelium during development. Nature 464, 108–111. 10.1038/nature08738.

39. Dunworth, W.P., Cardona-Costa, J., Bozkulak, E.C., Kim, J.D., Meadows, S., Fischer, J.C., Wang, Y., Cleaver, O., Qyang, Y., Ober, E.A., and Jin, S.W. (2014). Bone morphogenetic protein 2 signaling negatively modulates lymphatic development in vertebrate embryos. Circ Res 114, 56–66. 10.1161/CIRCRESAHA.114.302452.

40. Okuda, K.S., Astin, J.W., Misa, J.P., Flores, M.V., Crosier, K.E., and Crosier, P.S. (2012). lyve1 expression reveals novel lymphatic vessels and new mechanisms for lymphatic vessel development in zebrafish. Development 139, 2381–2391. 10.1242/dev.077701.

41. Asakawa, K., Suster, M.L., Mizusawa, K., Nagayoshi, S., Kotani, T., Urasaki, A., Kishimoto, Y., Hibi, M., and Kawakami, K. (2008). Genetic dissection of neural circuits by Tol2 transposon-mediated Gal4 gene and enhancer trapping in zebrafish. Proc Natl Acad Sci U S A 105, 1255–1260. 10.1073/pnas.0704963105.

42. Grzegorski, S.J., Chiari, E.F., Robbins, A., Kish, P.E., and Kahana, A. (2014). Natural variability of Kozak sequences correlates with function in a zebrafish model. PLoS One 9, e108475. 10.1371/journal.pone.0108475.

43. Diez, M., Medina-Munoz, S.G., Castellano, L.A., da Silva Pescador, G., Wu, Q., and Bazzini, A.A. (2022). iCodon customizes gene expression based on the codon composition. Sci Rep 12, 12126. 10.1038/s41598-022-15526-7.

44. Villefranc, J.A., Amigo, J., and Lawson, N.D. (2007). Gateway compatible vectors for analysis of gene function in the zebrafish. Dev Dyn 236, 3077–3087. 10.1002/dvdy.21354.

45. Choi, H.M.T., Schwarzkopf, M., Fornace, M.E., Acharya, A., Artavanis, G., Stegmaier, J., Cunha, A., and Pierce, N.A. (2018). Third-generation in situ hybridization chain reaction: multiplexed, quantitative, sensitive, versatile, robust. Development 145. 10.1242/dev.165753.

46. Jowett, T. (1999). Analysis of protein and gene expression. Methods Cell Biol 59, 63–85. 10.1016/s0091-679x(08)61821-x.

47. Reischauer, S., Stone, O.A., Villasenor, A., Chi, N., Jin, S.W., Martin, M., Lee, M.T., Fukuda, N., Marass, M., Witty, A., et al. (2016). Cloche is a bHLH-PAS transcription factor that drives haemato-vascular specification. Nature 535, 294–298. 10.1038/nature18614.

48. Sumanas, S., Jorniak, T., and Lin, S. (2005). Identification of novel vascular endothelial-specific genes by the microarray analysis of the zebrafish cloche mutants. Blood 106, 534–541. 10.1182/blood-2004-12-4653.

49. Liao, E.C., Paw, B.H., Oates, A.C., Pratt, S.J., Postlethwait, J.H., and Zon, L.I. (1998). SCL/Tal-1 transcription factor acts downstream of cloche to specify hematopoietic and vascular progenitors in zebrafish. Genes Dev 12, 621–626. 10.1101/gad.12.5.621.

50. Potter, A.S., and Steven Potter, S. (2019). Dissociation of Tissues for Single-Cell Analysis. Methods Mol Biol 1926, 55–62. 10.1007/978-1-4939-9021-4_5.

51. Hao, Y., Hao, S., Andersen-Nissen, E., Mauck, W.M., 3rd, Zheng, S., Butler, A., Lee, M.J., Wilk, A.J., Darby, C., Zager, M., et al. (2021). Integrated analysis of multimodal single-cell data. Cell 184, 3573–3587 e3529. 10.1016/j.cell.2021.04.048.

52. Wickham, H. (2016). ggplot2 : Elegant Graphics for Data Analysis. Use R!,. 2nd ed. Springer International Publishing : Imprint: Springer,.

53. Wickham H, A.M., Bryan J, Chang W, McGowan LD, François R, Grolemund G, Hayes A, Henry L, Hester J, Kuhn M, Pedersen TL, Miller E, Bache SM, Müller K, Ooms J, Robinson D, Seidel DP, Spinu V, Takahashi K, Vaughan D, Wilke C, Woo K, Yutani H (2019). Welcome to the tidyverse. Journal of Open Source Software. 10.21105/joss.01686.

54. Wickham, H. (2007). Reshaping Data with the reshape Package. Journal of Statistical Software 21(*12*), 1–20.

55. Wickham H, F.R., Henry L, Müller K, Vaughan D (2023). dplyr: A Grammar of Data Manipulation.

56. Hao, Y., Stuart, T., Kowalski, M.H., Choudhary, S., Hoffman, P., Hartman, A., Srivastava, A., Molla, G., Madad, S., Fernandez-Granda, C., and Satija, R. (2024). Dictionary learning for integrative, multimodal and scalable single-cell analysis. Nat Biotechnol 42, 293–304. 10.1038/s41587-023-01767-y.

57. R Core Team (2025). R: A language and environment for statistical computing.

58. Rstudio Team (2020). RStudio: Integrated Development for R (RStudio, Inc.).

